# Identification of a 2-substituted Benzimidazole Derivative with Antimycobacterial Activity and Synergistic Interaction with Rifampicin

**DOI:** 10.64898/2026.03.09.710429

**Authors:** Sarika Thakur, Alka Sharma, Narayan Kumar, Yangala Sudheer Babu, Mulaka Maruthi, Ravi Kumar, Ram Shankar Upadhayaya, Sumitra Nain, Ram Gopal Nitharwal

## Abstract

Infectious diseases remain a persistent global health burden, with bacterial infections accounting for a substantial disease burden. The increasing prevalence of antimicrobial resistance has emphasised the need for the development of novel antibacterial drugs. In the present study, the antibacterial activity of novel 2-substituted benzimidazole derivatives (NR-1 to NR-9) was evaluated against three bacteria, viz. *M. smegmatis, B. subtilis* and *E. coli* using estimation of Minimum Inhibitory Concentration (MIC) and Minimum Bactericidal Concentration (MBC). Two (NR-4 and NR-5) exhibited inhibitory activity against *M. smegmatis*, while two (NR-5 and NR-7) were active against *B. subtilis,* with MICs ranging from 62.5 to 250 μg/mL. Notably, NR-5 demonstrated antibacterial activity against both *M. smegmatis* and *B. subtilis*, with more efficacy against *M. smegmatis* (MICs: 125 μg/mL vs 62.5 μg/mL), bactericidal activity with an MBC/MIC ratio of 1 and low toxicity against Vero cells. Furthermore, NR-5 exhibited concentration-dependent killing kinetics and a synergistic interaction with Rifampicin (FICI: 0.498) against *M. smegmatis*. Molecular docking analysis predicted thymidylate kinase (Tmk) with a binding energy of -6.61 Kcal/mol as a plausible target of NR-5, and molecular dynamics simulation supported the stability of the Tmk-NR-5 complex over 100 ns of simulation. Thus, based on its potency, low-toxicity antimycobacterial activity and synergistic action with rifampicin, NR-5 is identified as a preliminary antimycobacterial lead against *M. smegmatis*.

## 1. Introduction

Infectious diseases have been a major contributor to mortality globally over the years. Several Gram-positive as well as Gram-negative bacteria, such as *Mycobacterium tuberculosis* (*M. tb*), *Bacillus cereus, Escherichia coli,* and *Staphylococcus aureus,* are responsible for several severe bacterial infections (Vianna et al. 2019; Amemiya et al. 2024). An acid-fast bacterium, *M. tb*, which is responsible for Tuberculosis (TB), one of the life-threatening infectious diseases, has a high prevalence rate. Other Gram-positive bacteria, such as *B. anthracis* and *B. cereus,* also cause deadly infections like anthrax and food-borne illness, respectively (Baindara and Aslam 2023). Since pathogenic strains are difficult to handle, surrogate models, such as *M. smegmatis* for *M. tb* (Lelovic et al. 2020)*, B. subtilis* for Gram-positive bacteria (Barák 2021; Stülke et al. 2023), and *E. coli* for Gram-negative bacteria (WHO 2018; Clinical and Laboratory Standards Institute 2023), are commonly used in laboratory research. Antibacterial drugs are essential tools for the treatment of bacterial infections, substantially reducing mortality and morbidity (Laxminarayan et al. 2016). These drugs kill or inhibit bacterial growth by targeting key cellular pathways, such as DNA replication, transcription, protein synthesis, and the cell wall (Murima et al. 2014; Baran et al. 2023; Thakur S et al. 2025). The conventional classes of antibiotics include penicillin, aminoglycosides, and fluoroquinolones, which are mainly used to treat bacterial infections. However, the improper use and overuse of existing antibiotics have led to the onset of antibiotic resistance, which has become a major global health threat (Sheikh et al. 2022). In present times, single-drug resistance and multiple-drug resistance have become defining characteristics of key pathogens. The World Health Organization (WHO) has highlighted antimicrobial resistance (AMR) as one of the top 10 health concerns globally (WHO 2024). The rising rate of resistance to existing drugs necessitates the discovery and development of novel antibacterial classes of drugs with novel mechanisms of action to overcome bacterial resistance (Miethke et al. 2021).

In this direction, heterocyclic compounds serve as promising pharmacologically relevant scaffolds in medicinal chemistry and serve as a core in several FDA-approved drugs (Pathak et al. 2026). Different classes such as triazoles, pyrazoles, benzimidazoles, thiophenes, oxadiazoles, etc., have been extensively studied in search of new antibacterial compounds (Bhatt et al. 2026; Patel et al. 2026; Pathak et al. 2026). Among these, benzimidazoles have become one of the preferred scaffolds for developing new inhibitory compounds in medicinal chemistry in recent years (Singh et al. 2012; Khan et al. 2024; Alheety et al. 2025). Benzimidazole derivatives display diverse antimicrobial properties, encompassing antifungal (Desai et al. 2014; El-Gohary and Shaaban 2017), antiviral (Xue et al. 2011; Pan et al. 2015), antibacterial (Mohanty et al. 2018), antimalarial (Morcoss et al. 2025),and antiparasitic (Anichina et al. 2021). Moreover, several clinically approved benzimidazole-derived drugs have been used for different purposes, such as antihelminthic, antiparasitic, and proton pump inhibitors for several years (Fig. 1). Although the benzimidazole scaffold is widely used in medicine, no clinically approved benzimidazole-derived antibacterial drug has been reported (Ibrahim et al. 2025; Tarek et al. 2025). One benzimidazole candidate, Ridinilazole, is currently in a phase III trial for the treatment of *Clostridioides difficile* (Okhuysen et al. 2024). Therefore, benzimidazole derivatives provide a new potential avenue for developing anti-resistance antibacterial agents. Benzimidazole is a heterocyclic compound that contains a benzene ring fused with an imidazole ring. It is considered a privileged scaffold due to its recurring appearance among various bioactive molecules. It can form various non-covalent intermolecular interactions, like metal ion coordination, π-π bonding, and hydrogen bonding, with several biological targets. The benzimidazole derivatives are synthesized by integrating different chemical groups, mainly at the 1-, 2-, and 5-positions of the benzimidazole ring. Also, the benzimidazole core has been postulated to serve as a purine analogue due to similarity in structure. Given that purines are important for the biosynthesis of nucleic acids and proteins in bacterial cells, benzimidazole derivatives may disrupt the synthesis of crucial components, leading to bacterial growth inhibition or cell death (Song and Ma 2016). Therefore, the designing of novel benzimidazole derivatives with different chemical groups is a prime need.

**Fig. 1.**
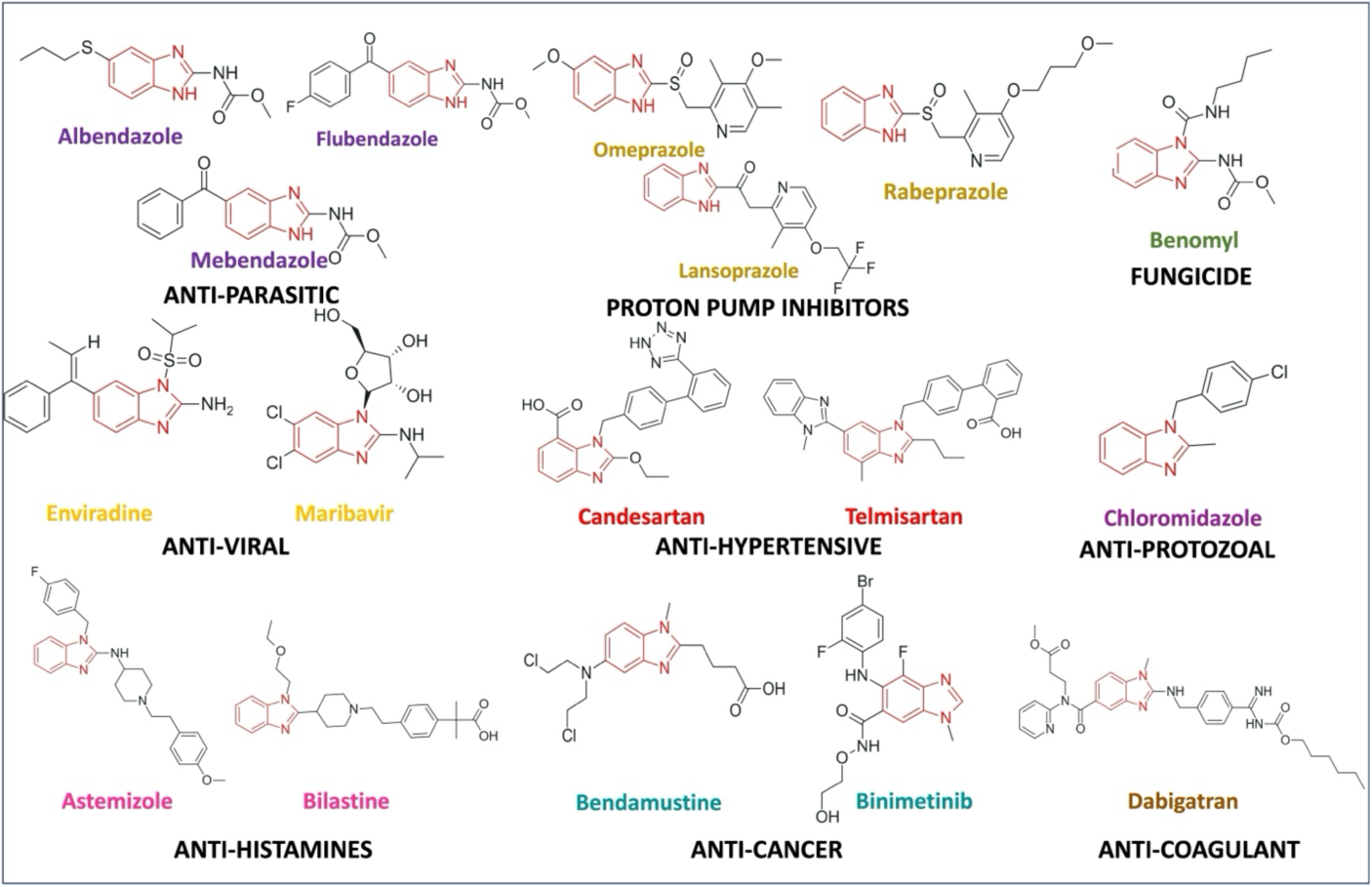
Benzimidazole-derived FDA-approved drugs. The benzimidazole core (highlighted in red) is present in several clinically approved drugs, which exhibit diverse inhibitory activities such as antiparasitic, antifungal, antiviral, and anticancer, etc. However, despite decades of extensive medicinal research efforts, there is no clinically approved antibacterial drug consisting of a benzimidazole scaffold to date.

In this study, we have evaluated the antibacterial activity of nine 2-substituted benzimidazole derivatives against three bacteria: *M. smegmatis*, *B. subtilis* and *E. coli*. These bacteria were used as the model system for acid-fast, gram-positive and Gram-negative bacteria, respectively. The antibacterial profiling of these compounds was done by various in vitro methods, including MIC, MBC, growth kinetics and time-kinetics evaluation. Furthermore, the cytotoxicity and ADME analysis were also performed to determine the drug-like properties of antibacterial benzimidazole compounds. The checkerboard assay was employed for the drug-drug interactions, and molecular docking and molecular dynamics simulation analysis were employed for in silico target prediction.

## 2. Methods

### 2.1. Benzimidazole Derivatives and Bacterial Strains

The small library of benzimidazole derivatives (NR1-NR9) was collected from the Pharmaceutical Chemistry Laboratory (Dr Sumitra Nain), Department of Pharmacy, Banasthali Vidyapeeth, Tonk, India (Chauhan et al. 2025) (Fig. 2). The stock solutions of benzimidazole derivatives were formulated in 100% sterile Dimethyl Sulfoxide (DMSO) (Hi-Media Ltd.). The antibacterial efficacy of the benzimidazole derivatives was determined against a panel of three bacteria: *Mycobacterium smegmatis mc^2^ 155*, *Bacillus subtilis MTCC121*, and *Escherichia coli ATCC25922*.

**Fig. 2.**
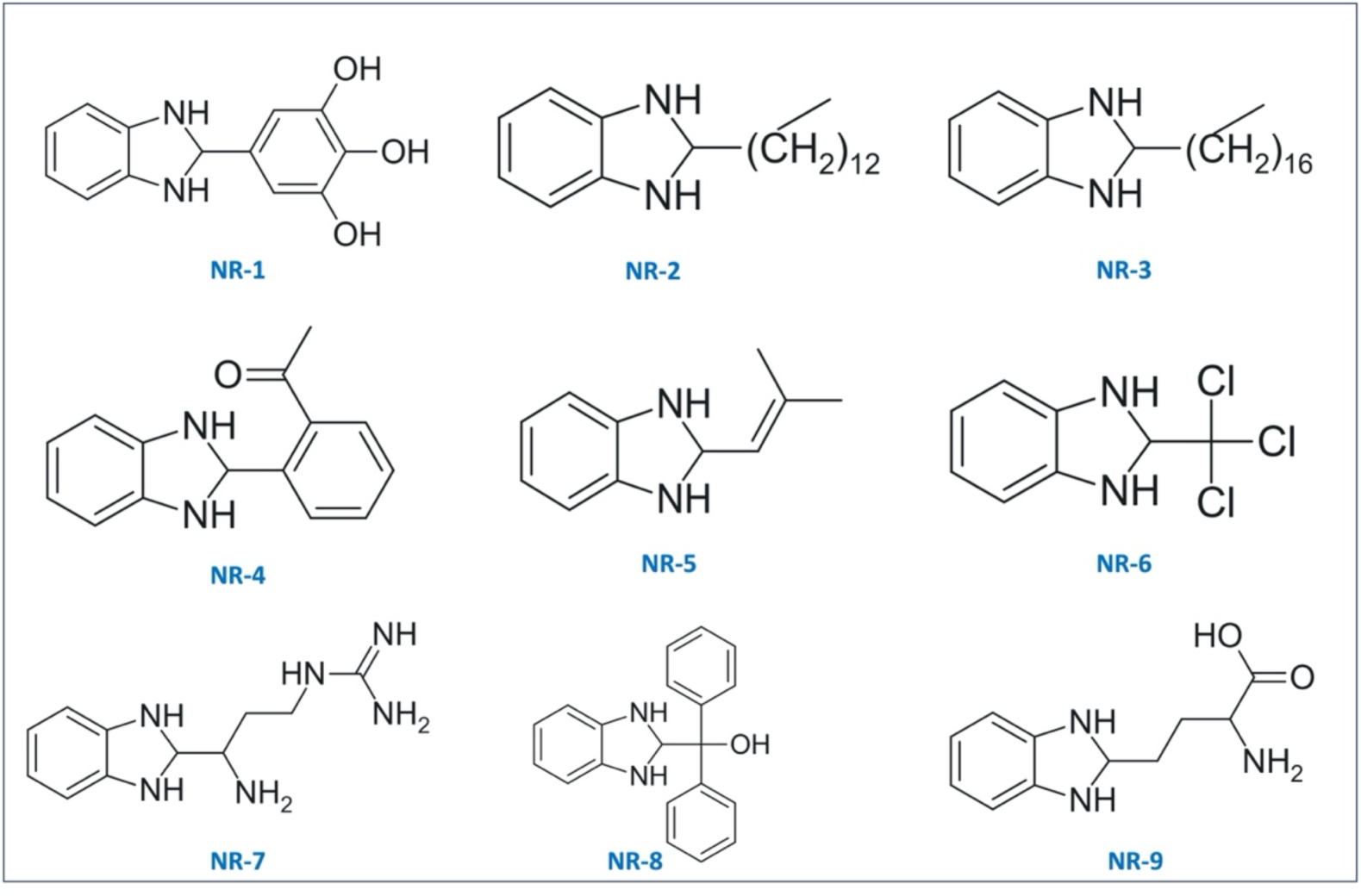
Chemical structures of the 2-substituted benzimidazole derivatives (NR-1-NR-9) used for antibacterial screening in this study.

### 2.2. Growth conditions of experimental strains

*M. smegmatis* was cultured on 7H9 Middlebrook medium supplemented with ADS (5g/L BSA, 2 g/L dextrose and 0.8 g/L NaCl) (Hi-Media Ltd) at 37℃ for 48 hours. *B. subtilis* and *E. coli* were grown on Mueller-Hinton (MH) broth and agar media and were incubated at 37℃ for 24 hours. The broth cultures were grown under shaking conditions (150 rpm) and agar cultures were grown under static conditions. For cytotoxicity analysis, a mammalian cell line obtained from the African green monkey’s kidney (epithelial cells), the Vero cell line, was cultured at 37℃ in Dulbecco’s Modified Eagle Medium (DMEM) supplemented with 10% Fetal Bovine Serum (FBS) and Penicillin-Streptomycin in a CO_2_ incubator.

### 2.3. Disc Diffusion assay

The disc diffusion assay was performed according to Clinical and Laboratory Standards Institute (CLSI) guidelines, with minor modifications for the preliminary evaluation (Clinical and Laboratory Standards Institute 2023). The cotton swabs were used to spread broth-grown cultures (O.D_600_-0.2) on the agar media plates (∼ 4 mm in depth) and allowed to dry for 15 minutes. Sterile discs (6 mm) were placed aseptically on agar Petri plates and spotted with 30 μL of benzimidazole derivatives (1 mg/mL), followed by a 30-minute diffusion period, then incubated at 37°C. Rifampicin (1 mg/mL) and Kanamycin (1 mg/mL) served as the positive control, while 10% DMSO (solvent) functioned as the negative control. Rifampicin was employed in this experiment because of its widely recognized anti-mycobacterial activity (White et al. 1971; Patel and Rosani 2023), and Kanamycin was selected because of its broad-spectrum antibacterial activity against Gram-positive and Gram-negative bacteria. The entire procedure was conducted using three independent biological replicates, with two technical duplicates in each experiment under a sterile environment, and zones of inhibition (ZOI) were measured after 24 hours of incubation.

### 2.4. Minimum Inhibitory Concentration (MIC) and Minimum Bactericidal Concentration (MBC) Determination

The minimum inhibitory concentration (MIC) of the susceptible derivative was measured using the Resazurin Microtitre Assay (REMA), with specific modifications (Banfi et al. 2003). A 96-well plate was used to assess the MIC, with two-fold serial dilutions of derivatives from 500 μg/mL to 7.81 μg/mL in growth media: 7H9 Middlebrook Broth for *M. smegmatis* and MH broth for *B. subtilis*. The inoculum was adjusted to approximately 5 × 10^5^ cfu/mL. Broth containing DMSO (starting at 5%) was used as a vector control, broth alone was employed as a sterility control, and broth containing inoculum was used as a growth control. 30 μL of 0.015% resazurin dye was added to each well after incubation at 37 °C for 48 hours for *M. smegmatis* and 24 hours for *B. subtilis*, and was further incubated overnight for *M. smegmatis* and for 4 hours for *B. subtilis*. The MIC was determined as the lowest concentration with no colour change from blue to pink. This experiment was performed using three independent biological replicates, each tested in technical duplicates. Following MIC determination, the MBC was assessed by the spot assay (Herigstad et al. 2001). The 10 μL culture from each well (up to and including the MIC value) from the 96-well MIC plate was spotted and cultured on the agar plate at 37℃ for 48 hours for *M. smegmatis* and 24 hours for *B. subtilis*. The concentration exhibiting no growth on the agar plate was reported as MBC. The MBC/MIC values were also estimated to determine the bactericidal or bacteriostatic mode of action of the derivatives.

### 2.5. Evaluation of the drug-likeliness of antibacterial benzimidazole derivatives

The Absorption, Distribution, Metabolism and Excretion (ADME) properties were evaluated *in* silico using SwissADME software (https://www.swissadme.ch/) (Daina et al. 2017). ADME key parameters involve Lipinski’s rule of five, a set of rules formed by Dr Christopher Lipinski, bioavailability, and gastrointestinal (GI) absorption. Lipinski’s rule of five is mainly used to evaluate whether a compound can be formulated into an orally active drug in humans (Jayavel et al. 2024). Moreover, toxicity prediction was done using ProTox 3.0 (Banerjee et al. 2024), following approaches reported in previous studies (Nepolraj et al. 2021; Jayavel et al. 2024). The analysis included toxicity class, hepatotoxicity, carcinogenicity, immunotoxicity, mutagenicity, and cytotoxicity predictions.

### 2.6. Cytotoxicity analysis

The cytotoxicity effect of the benzimidazole derivatives was assessed on the Vero cell line using the MTT (3-(4,5-dimethylthiazol-2-yl)-2, 5-diphenyl tetrazolium bromide) assay. The viability of cells was determined using yellow (3-(4,5-dimethylthiazole-2-yl)-2,5-biphenyl tetrazolium bromide (MTT) salt solution [5 mg/ml in 1X Phosphate Buffer Saline (PBS)]. The Vero cells were collected by trypsinization (0.25% trypsin-EDTA). In a 96-well plate, 1 × 10^4^ cells/well were seeded and incubated in the CO_2_ incubator for 24 hours for cell growth and attachment to the wells. After 24 hours, test compounds with varying concentrations (500 μg/mL as the highest concentration) were added to the 96-well plate and incubated for 48 hours. The MTT salt solution was added to the wells after the compounds were washed off with 1X PBS. The culture plate was incubated at 37℃ for 3 hours. In MTT assays, all the appropriate controls were used, including a vehicle control. Untreated cells in wells were considered as the negative control; cell-free wells were used as blanks. Although the compounds were washed off gently before addition of MTT, compound-related interference with MTT was not assessed separately. Our final absorbance was calculated after subtraction of the experimental background vehicle control at 570nm. After incubation, DMSO was added to dissolve the formazan crystals, and after 30 minutes of incubation, the absorbance was measured at 570 nm. IC_50_ values for each test compound were analyzed in GraphPad Prism 8.0.1 by fitting the results to a sigmoidal equation. The selectivity Index (SI) was evaluated as a ratio of IC_50_ against the Vero cell line and MIC against the bacteria.

### 2.7. Effect of NR-5 on the Growth Kinetics of *M. smegmatis*

A growth-kinetics assay of NR-5 was conducted using three different concentrations, 0.5x, 1x and 2x of MIC. Broth consisting of the derivative at different concentrations (0.5x MIC, 1x MIC and 2x MIC), a positive control and a growth control was inoculated with the primary culture, so that the starting OD_600_ was 0.01. The 500 μL aliquots were collected at different time intervals (0, 12, 24, 36, and 48 hours), and the optical density at 600 nm was monitored. Rifampicin (1x MIC) was used as a positive control, which acted as a reference drug for antimycobacterial activity. The experiment was performed in three independent biological replicates (N=3). We assessed *M. smegmatis* growth kinetics by fitting the Gompertz growth model and performed statistical analyses to test significance (p < 0.5) using GraphPad Prism 8.0.1. The model yielded the following parameters for each treatment: maximum asymptotic optical density (Ym), growth rate constant (K), initial optical density (Y_o_). Goodness of fit was estimated by the coefficient of determination (R^2^).

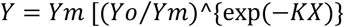

Where Y_o_ is the starting population, Y_m_ is the maximum population, and K is the growth rate constant that describes the lag time. A time vs absorbance curve was constructed to evaluate the effect of the derivative on the growth of *M. smegmatis*.

### 2.8. Kill kinetics of NR-5 against *M. smegmatis* under non-growing conditions

*M. smegmatis* was cultured and grown at 37℃ till exponential phase (0.4-0.6). The cells were centrifuged at 12000 rpm for 10 minutes. Then the cell pellet was washed twice with 1X PBS and resuspended in PBS. After this, the cells were adjusted to 5 × 10^5^ CFU/mL in PBS, followed by treatment with a 2x MIC concentration of NR-5. The cells were incubated at 37℃, and then 100 μL of 10^-2^ dilution was plated on 7H9 agar plates at different time intervals (0, 2, 4, 6 and 8 hours). The agar plates were incubated at 37℃ for 48 hours, and then the colony count was done. The experiment was performed independently three times. Further, to minimize residual compound carryover, the treated bacterial suspensions were serially diluted to 100 times before plating to reduce the concentration of derivatives.

### 2.9. Analysis of the drug-drug interaction of benzimidazole derivative NR-5 with clinically approved first- and second-line anti-TB drugs

Initially, the MIC values of the susceptible antibiotics, rifampicin, ethambutol, moxifloxacin, kanamycin, streptomycin and bedaquiline were determined against *M. smegmatis* by REMA as described earlier, followed by the checkerboard assay to determine the drug-drug interactions. The checkerboard assay was performed to assess the synergistic interaction of the most effective benzimidazole derivative, NR-5, with clinically approved first- and second-line anti-TB drugs against *M. smegmatis*. This assay was performed using a 96-well plate resazurin assay as described previously, with a few modifications (Bellio et al. 2021). It was performed three times independently (N=3) to ensure reproducibility of the observed interaction patterns. The fractional inhibitory concentration index (FICI) was calculated using the index equation described by Berenbaum (Berenbaum 1977).

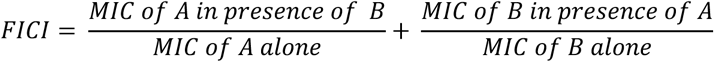

A FICI value of ≤ 0.50 was defined as synergism, 0.51-1 as additive, 1-4 as indifferent and >4 as antagonism (Doern 2014). After the checkerboard analysis, the effect of the synergistic combination concentration, the lowest concentration at which NR-5 inhibits growth in combination, on the growth kinetics of *M. smegmatis* was also assessed.

### 2.10. In silico Molecular Docking Analysis of NR-5 with essential proteins of *M. smegmatis*

Molecular docking analysis was done to find a plausible target for the derivative NR-5. The structures of 460 essential proteins of *M. smegmatis* (DeJesus et al. 2017; Dragset et al. 2019) were downloaded from the Protein Data Bank (PDB) and the AlphaFold database using the structure identifiers in the PDB format (Supplementary datasheet). For an AlphaFold-derived structure, model confidence was evaluated using the AlphaFold predicted Local Distance Difference Test (pLDDT) scores (pLDDT > 70). Since the study comprised a large-scale screening against 460 protein targets, individual pLDDT values are not presented. The 2D structure of the ligand (NR-5) was drawn in Marvin Sketch (ChemAxon) and converted to a 3D structure in PDB format in Avogadro 1.2.0, followed by optimization. The protein structures were prepared for docking studies by removing irrelevant residues and ligands through Discovery Studio Visualizer. Co-crystallized heteroatoms and water molecules were excluded, and polar hydrogens and Kollman charges were added in the docking workflow. Docking analysis of NR-5 was performed against 460 protein targets; therefore, the grid centre and dimensions were adjusted according to the spatial coordinates and dimensions of each protein model. Molecular docking was performed using AutoDock version 4.2.6 following the Lamarckian genetic algorithm protocol. For each docked protein-ligand complex, several docking conformations were obtained, and the conformation with the least binding energy was selected for further analysis. The protein binding pockets were predicted using the PrankWeb server (Polák et al. 2025), followed by redocking with AutoDock Vina. Protein-ligand interactions, such as hydrogen bonding and hydrophobic interactions, were investigated and visualised using Discovery Studio Visualizer 2025 and PyMOL.

### 2.11. MD Simulation

Molecular dynamics (MD) simulations were performed using GROMACS 2023.3 (Abraham et al. 2015) to evaluate the stability and dynamic behaviour of the NR-5-thymidylate kinase (Tmk) complex. The protein structure was prepared by removing water molecules and adding missing hydrogen atoms. The CHARMM36m force field was used for the protein, while the ligand parameters were generated using the CGenFF server (Vanommeslaeghe et al. 2010). The complex was placed in a cubic box with a minimum 10 Å distance from the box edges, solvated with the SPC/E water model, and neutralized with counterions. NaCl was added to maintain a physiological ionic concentration of 0.15 M.

The system was first energy-minimized using the steepest descent algorithm and then equilibrated at 300 K under the NVT ensemble for 100 ps using the V-rescale thermostat. Hydrogen bonds were constrained using LINCS, and long-range electrostatic interactions were treated using the Particle Mesh Ewald (PME) method with a 1.2 nm cutoff.

A 100 ns production MD simulation was performed as ten consecutive 10 ns runs, with each run continued from the checkpoint file of the previous segment. The resulting trajectories were combined into a continuous trajectory, followed by periodic boundary correction, centring, and structural alignment. The final trajectory was analyzed for RMSD, RMSF, radius of gyration (Rg), hydrogen bonding, and minimum distance. The analysis plots were generated using Python-based scripts.

## 3. Results

### 3.1. Identification of antibacterial benzimidazole derivatives

In an effort to identify novel antibacterials, a small library of nine recently synthesized benzimidazole derivatives (NR1-NR9) (Chauhan et al. 2025) was evaluated for antibacterial activity against a panel of three bacteria representing acid-fast (*M. smegmatis*), Gram-positive (*B. subtilis*) and Gram-negative (*E. coli*). Preliminary phenotypic screening for antibacterial activity was performed using the disc diffusion assay. The antifungal potential of these compounds against *Candida tropicalis* has been previously reported (Chauhan et al. 2025). Among the nine derivatives (NR1-NR9) tested, three derivatives, NR-4, NR-5, and NR-7, exhibited promising antibacterial activity with significant zones of inhibition (ZOI) (≥10 mm) at a concentration of 1 mg/mL (Fig. 3). Specifically, NR-4 and NR-5 exhibited antimycobacterial activity against *M. smegmatis*, whereas NR-5 and NR-7 showed inhibitory activity against *B. subtilis*. None of the nine derivatives screened showed inhibitory activity against *E. coli*. Notably, NR-5 displayed a zone of inhibition comparable to rifampicin against *M. smegmatis*.

**Fig. 3.**
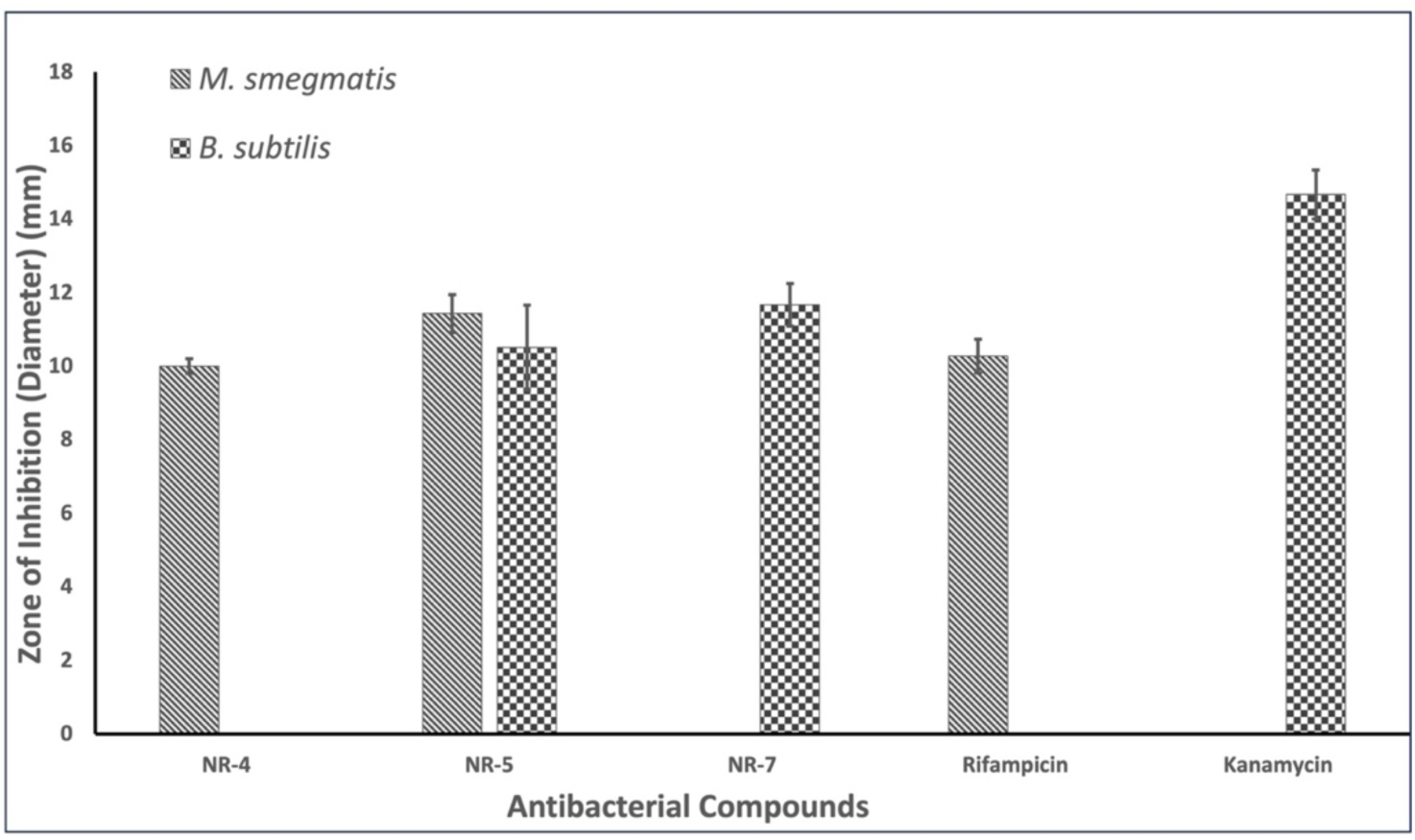
Preliminary assessment of antibacterial activity of benzimidazole derivatives. The zone of inhibition was estimated using the disc diffusion assay against a panel of three bacteria, *M. smegmatis*, *B. subtilis, and E. coli,* representing acid-fast, Gram-positive, and Gram-negative bacteria. None of the compounds showed activity against *E. coli*.

### 3.2. Determination of antibacterial activity of Benzimidazole derivatives

To further characterize the compounds that demonstrated activity during the primary screening, namely NR-4, NR-5, and NR-7, their minimum inhibitory concentrations (MICs) were determined using the resazurin microdilution assay (Table 1). Among the tested derivatives, NR-5 emerged as the most efficient derivative with an MIC of 62.5 μg/mL, followed by NR-4 with an MIC of 250 μg/mL against *M. smegmatis.* (Table 1). Under the experimental conditions used in this study, rifampicin exhibited an MIC of 31.25 μg/mL (against *M. smegmatis*), suggesting that NR-5 possesses substantial antimycobacterial activity, although it is approximately two-fold less potent than the reference drug. Against *B. subtilis,* both the NR-5 and NR-7 derivatives exhibited an MIC value of 125 μg/mL.

**Table 1.** In vitro antibacterial profiling of benzimidazole derivatives to determine the mode of antibacterial action.

| Benzimidazole derivatives | *Mycobacterium smegmatis* |  |  |  | *Bacillus subtilis* |  |  |  |
| --- | --- | --- | --- | --- | --- | --- | --- | --- |
|  | MIC (µg/mL) | MBC (µg/mL) | MBC/MIC Ratio<sup>a</sup> | Type of activity | MIC (µg/mL) | MBC (µg/mL) | MBC/MIC Ratio<sup>a</sup> | Type of activity |
| NR-4 | 250 | 500 | 2 | Bactericidal | - | - | - |  |
| NR-5 | 62.5 | 62.5 | 1 | Bactericidal | 125 | 250 | 2 | Bactericidal |
| NR-7 | - | - | - | - | 125 | 250 | 2 | Bactericidal |
| Rifampicin | 31.25 | 62.5 | 2 | Bactericidal | - | - | - | - |
| Kanamycin | - | - | - | - | 7.8 | 31.25 | 4 | Bactericidal |

<sup>a</sup>MBC/MIC ratio: MBC/MIC $\leq 4$ = bactericidal; $> 4$ = bacteriostatic.

In addition to inhibitory activity, it is important to determine whether an antibacterial derivative exerts a bacteriostatic or bactericidal effect. One widely accepted approach is the evaluation of the minimum bactericidal concentration (MBC) to MIC ratio. The MBC/MIC ratio provides insights into the killing potency of a compound and, thus, is an important parameter in drug development strategies. Thereby, the MBC values of active compounds were determined, and their MBC/MIC ratio were calculated. (Table 1). The derivatives are defined as bactericidal if the MBC/MIC ratio is ≤ 4, whereas a value > 4 indicates a bacteriostatic effect (Ishak et al. 2025). The analysis of the MBC/MIC ratios revealed that NR-4, NR-5, and NR-7 exert bactericidal activity against their specific susceptible bacterial strains (Table 1). Importantly, NR-5 displayed the lowest MIC value against *M. smegmatis* (62.5 μg/mL) as well as the lowest MBC/MIC ratio, highlighting its pronounced antimycobacterial efficacy among the tested derivatives (Fig. 4).

**Fig. 4.**
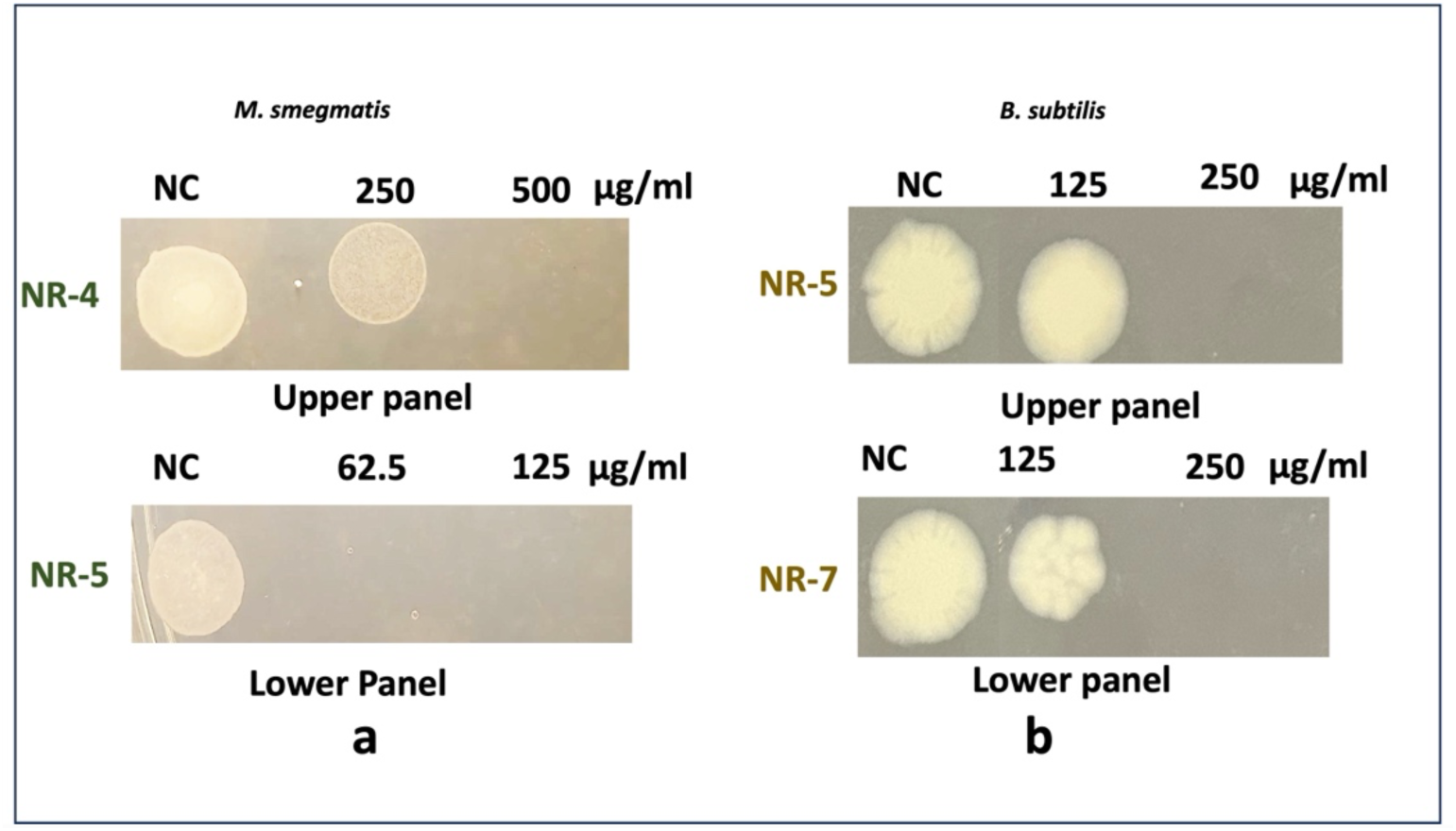
Determination of Minimum Bactericidal Concentration (MBC) of Benzimidazole Derivatives against *M. smegmatis* and *B. subtilis*. The MBC of the benzimidazole derivatives (NR-4, NR-5 and NR-7) was determined using the drop dilution method and the concentration with no visible growth on the agar was reported as the MBC. The derivative NR-5 displayed better bactericidal activity than the other antibacterial derivatives (NR4 and NR-7), highlighting its strong antibacterial action against *M. smegmatis*.

### 3.3. Prediction of drug-likeness of antibacterial Benzimidazole derivatives

Based on the MIC and MBC analyses, three benzimidazole derivatives—NR-4, NR-5, and NR-7—were identified as the promising antibacterial candidates and were therefore subjected to in silico ADME (Absorption, Distribution, Metabolism, and Excretion) evaluation using the SwissADME platform. Prediction of the pharmacokinetic profile of a compound is a critical step in the drug discovery process. Early assessment of physicochemical and pharmacokinetic properties facilitates the identification of compounds with favourable drug-like characteristics, thereby reducing the likelihood of failure during later stages of drug development and improving the probability of successful target engagement.

The present analysis was conducted to evaluate the drug-likeness and predict the pharmacokinetic behaviour of the active benzimidazole derivatives. SwissADME bioavailability radar plots (Fig. 5) revealed that all the antibacterial benzimidazole derivatives displayed physicochemical properties within or close to the optimal range for oral bioavailability based on lipophilicity (−0.7 to +5), molecular weight (150 to 500 g/mol), polarity (−20 to 130 Å^2^), solubility (<6), saturation (>0.25), and flexibility (<9 rotational bonds) (Table 2). Also, the ADME prediction revealed that compound NR-5 exhibited high gastrointestinal absorption. Though it was predicted that the compound possesses blood-brain barrier permeability (Supplementary Fig. 1), it was not considered an exclusion criterion due to its overall drug-likeness and pharmacokinetic profile (Supplementary Table 4). The BBB permeability may facilitate CNS exposure and therefore needs further evaluation in future pharmacokinetic and safety studies.

**Fig. 5.**
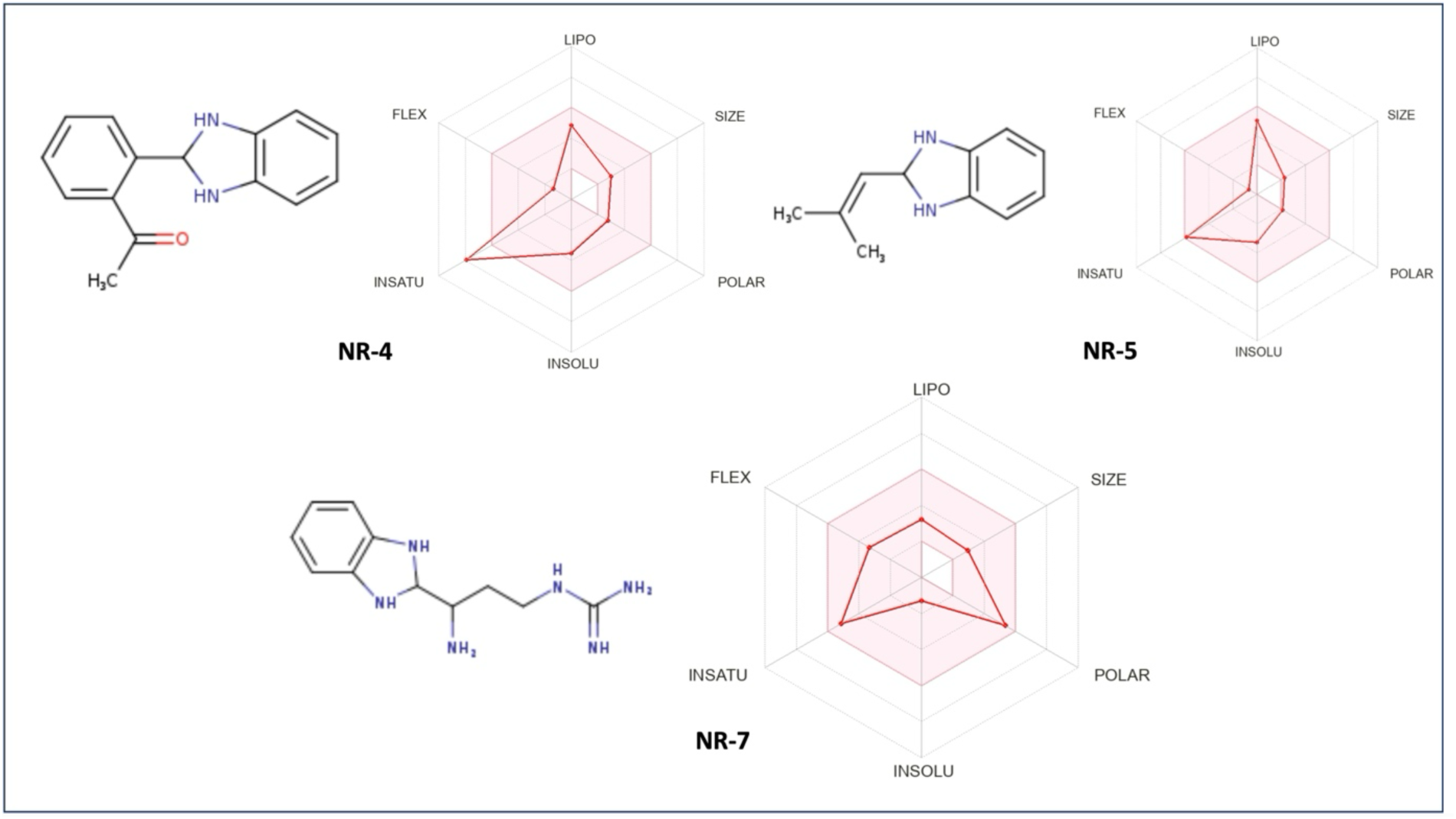
SwissADME bioavailability radar plots of antibacterial benzimidazole derivatives NR-4, NR-5 and NR-7. Each radar depicts the oral bioavailability of the derivative, where the pink region highlights the optimal physicochemical range in terms of lipophilicity, polarity, molecular size, solubility, saturation and flexibility. It was observed that two antibacterial derivatives (NR-5 and NR-7) displayed values within the optimal range of physicochemical parameters; however, NR-4 failed to meet the recommended saturation value ( > 0.25), suggesting the structural optimization of the derivative.

**Table 2.** Physicochemical Properties of 2-substituted Benzimidazole Derivatives.

| S.No. | Parameters | NR-4 | NR-5 | NR-7 |
| --- | --- | --- | --- | --- |
| 1 | Lipophilicity (XlogP3) | 2.97 | 3.30 | 0.14 |
| 2 | Molecular weight (g/mol) | 238.28 | 174.24 | 234.30 |
| 3 | Polarity (TPSA) ($\text{\AA}^2$) | 41.13 | 24.06 | 111.98 |
| 4 | Solubility (logS) | -3.55 | -3.27 | -1.31 |
| 5 | Saturation (sp<sup>3</sup> hybridization) | 0.13 | 0.27 | 0.36 |
| 6 | Flexibility (Rotatable bonds) | 2 | 1 | 5 |

The *in silico* toxicity profiles of NR-4, NR-5, and NR-7 were evaluated using the ProTox 3.0 web server. NR-5 exhibited a comparatively favourable predicted profile as it displayed inactivity for hepatotoxicity, carcinogenicity, immunotoxicity, mutagenicity, and cytotoxicity, whereas NR-4 was predicted to be active for hepatotoxicity, carcinogenicity, and mutagenicity. NR-7 also showed inactive predictions for these five endpoints (Supplementary Table 5). Collectively, NR-5 exhibited a less toxicological profile. These findings are based solely on computational predictions, and further experimental investigations are needed to confirm its safety profile.

### 3.4. Cytotoxicity and selectivity assessment of antibacterial Benzimidazole Derivatives in Vero Cells

The cytotoxic potential of the three most active antibacterial benzimidazole derivatives, NR-4, NR-5 and NR-7, was evaluated against Vero kidney epithelial cells using the MTT assay. The compounds were tested at concentrations up to 500 μg/mL, with rifampicin serving as the reference drug. Among the tested derivatives, NR-4 and NR-5 exhibited comparatively low cytotoxicity toward Vero cells, with IC_50_ values of 170.65 ± 6.49 μg/mL and 178.35 ± 9.18 μg/mL, respectively (Table 3). These values were considerably higher than rifampicin (IC_50_ = 41.51 ± 6.00 μg/mL), revealing lower toxicity to mammalian cells compared to rifampicin. However, NR-7 exhibited a higher cytotoxic effect (IC_50_ = 32.60 ± 4.46 μg/mL), which was slightly lower than that of rifampicin (Table 3). The current study is a preliminary assessment of cytotoxicity shown by the compounds in Vero cells. NR-5 showed limited cytotoxicity in Vero cells with moderate selectivity (SI) as an antibacterial, where experiments were performed in triplicate (n=3) over three independent days. Statistical analysis for NR-4, NR-5, NR-7, and Rifampicin (Fig. 6) was performed using one-way ANOVA with comparisons between controls and compounds, and statistically significant differences were obtained (p < 0.0001 ****; p-value summary, p < 0.05; Fig. 6).

**Table 3.** Cytotoxic efficacy and Selectivity Index of compounds along with the standard drug (rifampicin), inhibitory concentrations (IC_50_) on Vero cells in μg/mL. N=3 (Number of experiments performed). The IC_50_ results were represented as the mean ± standard error of the mean (S.E.M.).

| S. No. | Compounds | $IC_{50}$ values ($\mu\text{g/mL}$) | MIC ($\mu\text{g/mL}$) | | Selectivity Index (SI) |
| --- | --- | --- | --- | --- | --- |
|  |  |  | *M. smegmatis* | *B. subtilis* |  |
| 1 | NR-4 | $170.65 \pm 6.49$ | 250 | - | 0.68 |
| 2 | NR-5 | $178.35 \pm 9.18$ | 62.5 | 125 | 2.85 |
| 3 | NR-7 | $32.6 \pm 4.46$ | - | 125 | 0.26 |
| 4 | Rifampicin | $41.51 \pm 6.0$ | 31.25 | - | 1.32 |

**Fig. 6.**
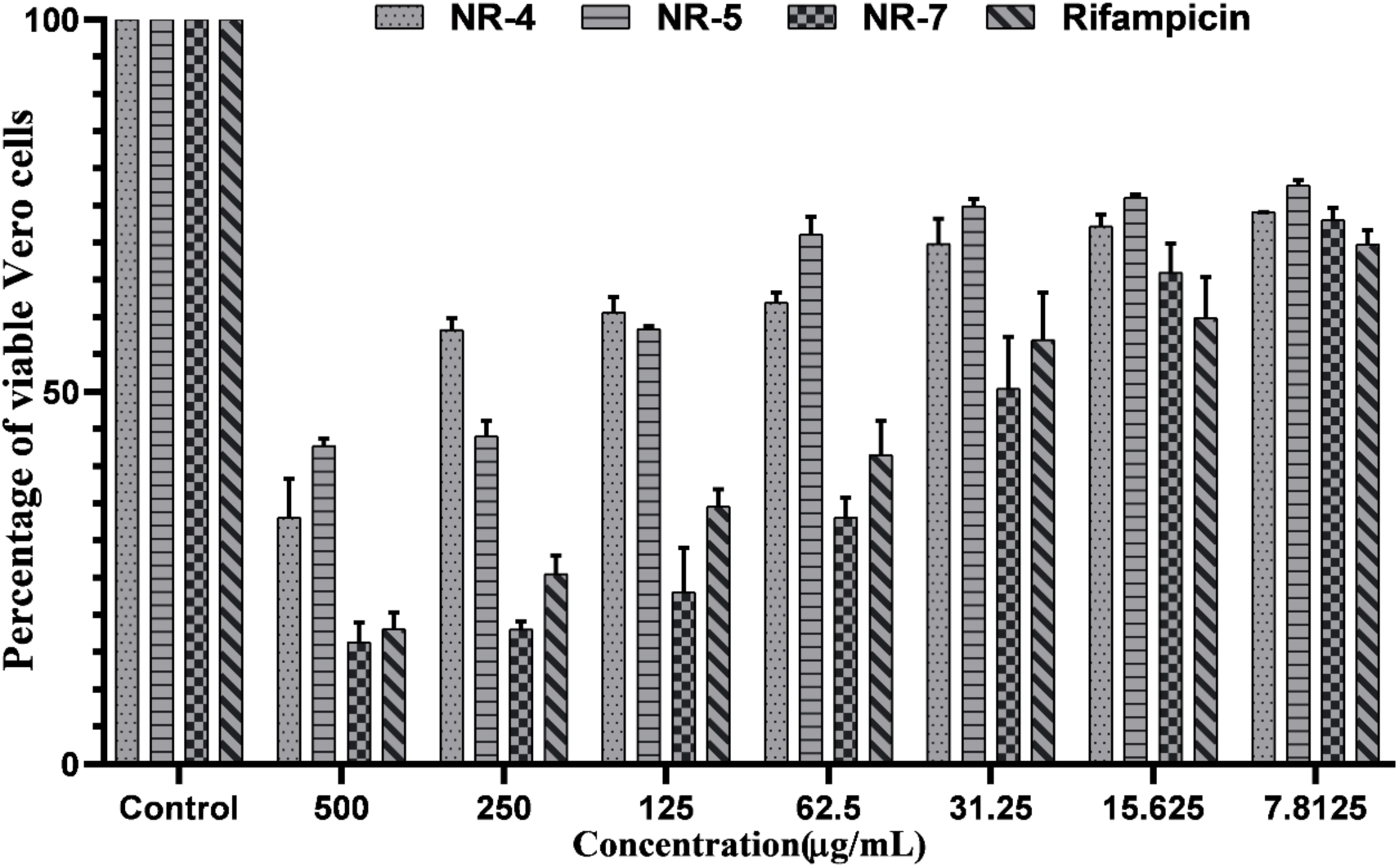
Cell viability analysis of benzimidazole derivatives in Vero cells. The percentage of viable Vero cells after treatment with varying concentrations of compounds NR-4, NR-5, and NR-7 was determined using a colourimetric assay (MTT assay). Rifampicin was used as a standard drug for control. The tested compounds displayed a concentration-dependent reduction in the viability of cells. Among the tested compounds, NR-5 exhibited the lowest cytotoxic effect, suggesting a favourable safety profile for further biological assessment.

The selectivity index (SI), calculated as the ratio of IC_50_ to MIC, was used to assess the therapeutic selectivity of the compounds. Among the tested derivatives, NR-5 exhibited the highest SI value (2.85), exceeding that of rifampicin (1.32), indicating greater selectivity toward mycobacterial cells over mammalian cells. However, NR-4 and NR-7 displayed poor selectivity with lower SI values of 0.68 and 0.26, respectively (Table 3). These findings cumulatively highlight that NR-5 possesses the favourable balance between antimycobacterial activity and cytotoxicity, which highlights NR-5 as a promising derivative for requiring further evaluations in order to develop this into lead molecule for antimycobacterial drug discovery.

### 3.5. Growth kinetics analysis reveals the concentration-dependent growth inhibitory activity of NR-5 against *M. smegmatis*

Since NR-5 emerged as the most potent antibacterial benzimidazole derivative based on antibacterial screening and safety profiling, its effect on the growth of *M. smegmatis* was further evaluated through growth kinetics analysis at different concentrations (0.5x, 1x, and 2x MIC). Growth kinetics were evaluated using the Gompertz model, a sigmoidal growth model. This model is extensively employed to describe microbial growth dynamics (Zwietering et al. 1990; Garcia et al. 2009; Hossain et al. 2016; Wawrzyniak 2020). The results revealed that the compound NR-5 displayed a concentration-dependent inhibitory activity on bacterial growth, resulting in a marked reduction in growth compared with the untreated control. Treatment with NR-5 at 0.5x and 1x MIC significantly suppressed bacterial growth, whereas complete growth inhibition was observed at 2x MIC (Fig. 7). The initial OD_600_ of the NR-5 treatments was experimentally set to 0.01 for all samples. The different values reported (0.0017, 0.002, and 0.011) were the initial values estimated by the Gompertz model during curve fitting and not the experimentally measured starting values. The Gompertz model adequately described the growth profiles of both treated and untreated cultures, as indicated by the high goodness-of-fit (R²) values obtained for all datasets. Moreover, Two-way repeated-measures ANOVA revealed a statistically significant effect of treatment (p < 0.0001), time (p < 0.0001), and their interaction (p < 0.0001) (Supplementary Table 6). This highlights that the bacterial growth profiles differed significantly among treatments over the incubation period.

**Fig. 7.**
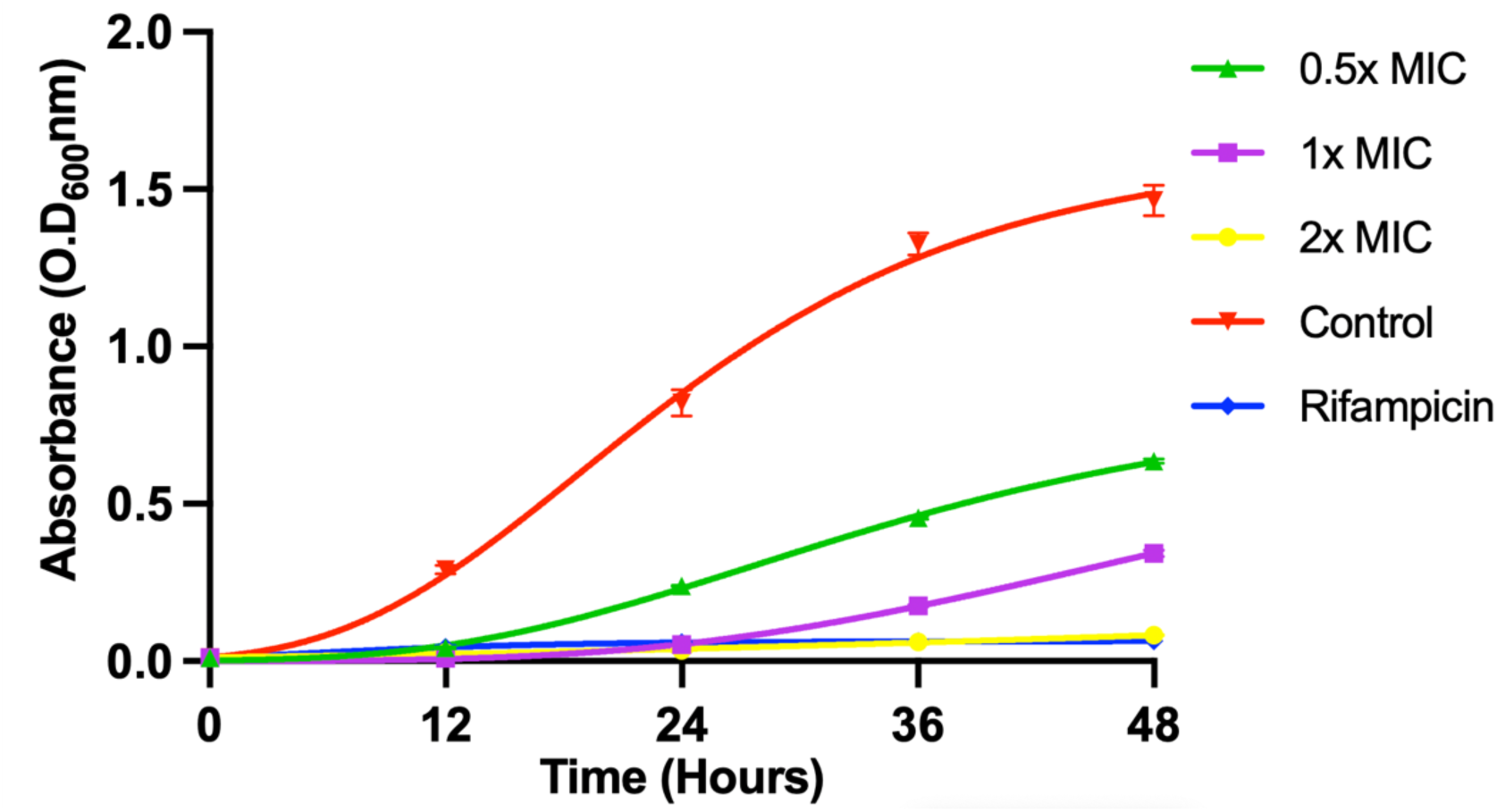
Growth kinetics profiles of *M. smegmatis* treated with NR-5 at different concentrations (0.5× MIC, 1× MIC, and 2× MIC) compared with the untreated negative control (no drug) and the positive control (rifampicin). The growth curves demonstrate a concentration-dependent inhibitory effect of NR-5 on bacterial growth, with progressively greater suppression observed at increasing concentrations. Complete growth inhibition was achieved at 2× MIC. Growth kinetics analysis confirmed the effective antibacterial activity of NR-5 against *M. smegmatis*. These findings highlight the strong growth-inhibitory activity of NR-5 and support its further evaluation as a potential antimycobacterial lead compound for further optimization.

Quantitative analysis showed that NR-5 reduced the bacterial growth by ∼2-fold, 4-fold and 7-fold at 0.5x, 1x, and 2x MIC, respectively. Furthermore, a progressive decrease in the growth rate constant was observed with increasing concentrations of NR-5, indicating a concentration-dependent reduction in bacterial growth potential (Table 4). The R^2^ value validates the model fitting and also the reliability of the growth kinetics parameters (Table 4). All these parameters together indicate that NR-5 has a strong inhibitory effect against *M. smegmatis*.

**Table 4.** Key parameters of the Gompertz model, employed to evaluate the effect of NR-5 on the growth kinetics of *M. smegmatis*.

| S.No. | Parameters | NR-5<br>(0.5x MIC) | NR-5<br>(1x MIC) | NR-5<br>(2x MIC) | Negative<br>control | Positive<br>control<br>(Rifampicin) |
| --- | --- | --- | --- | --- | --- | --- |
| 1 | Y<sub>m</sub><br>(Maximum growth) | 0.98 | 0.85 | 0.31 | 1.615 | 0.063 |
| 2 | Y<sub>0</sub> (Initial OD) | 0.0017 | 0.002 | 0.011 | 0.011 | 0.011 |
| 3 | K (Growth rate<br>constant) | 0.065 | 0.046 | 0.018 | 0.084 | 0.13 |
| 4 | R<sup>2</sup> (Coefficient of<br>determination) | 0.99 | 0.99 | 0.97 | 0.99 | 0.98 |

### 3.6. Time-kill kinetics confirm the rapid bactericidal effect of NR-5 against *M. smegmatis*

The growth kinetics analysis revealed that NR-5 significantly inhibits the growth of *M. smegmatis*, indicating its efficacy against actively growing cells. To determine whether the antibacterial activity of NR-5 extends beyond growth inhibition, time-kill kinetics assays were conducted under non-growing conditions. The treatment with NR-5 at 2x MIC concentration was given to *M. smegmatis* suspended in 1× PBS (5 × 10^5^ cfu/mL), which resulted in a progressive reduction in CFU/mL (Colony Forming Unit/mL) over time compared with the untreated control (Fig. 8 (b)). Notably, after 8 hours of exposure, NR-5 completely killed the bacteria, as evidenced by the absence of detectable colonies on the 7H9 agar plates following 48 hours of incubation (Fig. 8a). This highlights the anti-mycobacterial activity of NR-5 with a ∼3-log reduction in viable bacterial counts.

**Fig. 8.**
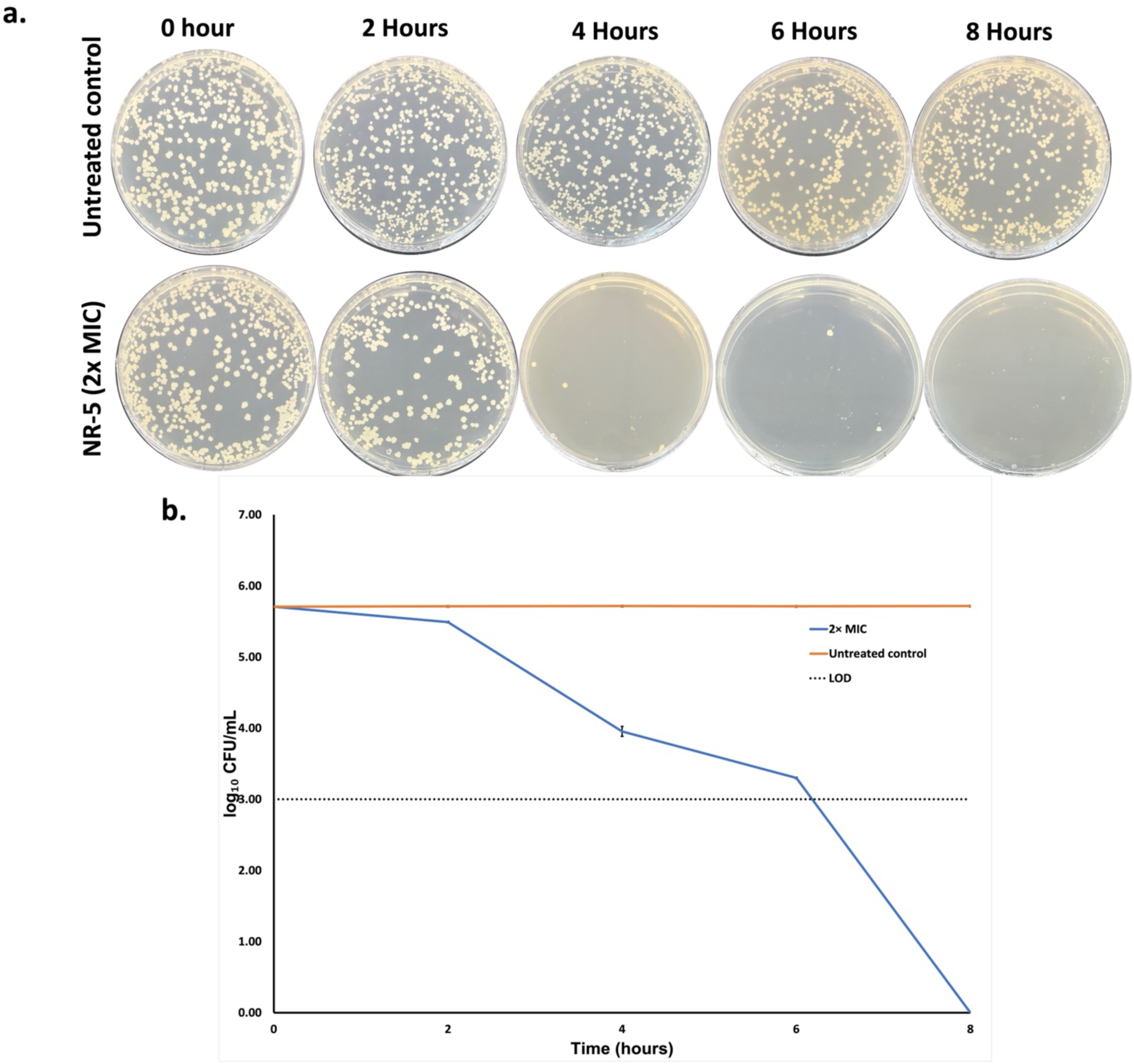
Kill Kinetics analysis of NR-5 against *M. smegmatis* under non-growing conditions. **(a)** Representative 7H9 agar plates showing a marked reduction in *M. smegmatis* colonies following treatment with NR-5 (2× MIC) under non-growing conditions. Fewer than 10 colonies were observed after 4 h of treatment compared with the untreated control, while no detectable viable colonies were observed after 8 h of exposure, following 48 h of incubation. **(b)** Time-kill kinetics curve depicting the reduction in viable bacterial counts (log₁₀ CFU/mL) of *M. smegmatis* treated with NR-5 at 2× MIC over an 8 h period. NR-5 induced a progressive decline in bacterial viability (∼ 3-log reduction), thereby confirming its bactericidal activity. The theoretical limit of detection (LOD) of the viable-count assay was 1 × 10³ CFU/mL (3 log₁₀ CFU/mL), based on plating 100 µL of a 10⁻² dilution. Samples yielding no detectable colonies were considered below the detection limit.

### 3.7. NR-5 displays synergistic interaction with rifampicin but indifferent interactions with other clinically approved anti-TB Drugs assessed against *M. smegmatis*

The two-dimensional checkerboard assay was employed to investigate the synergistic interaction between NR-5 and a panel of anti-TB drugs of different classes and mechanisms of action against *M. smegmatis*. Drug interactions were evaluated by the estimation of the Fractional Inhibitory Concentration Index (FICI) (see methods section).

Among the combinations tested, NR-5 displayed synergistic interaction with rifampicin. When used in combination, the minimum inhibitory concentrations (MICs) of both NR-5 and rifampicin were reduced fourfold compared with their respective MIC values when used alone (Fig. 9). This synergistic effect was further supported by a FICI value of 0.498, indicating enhanced antimycobacterial activity of the combination (Table 5).

**Fig. 9.**
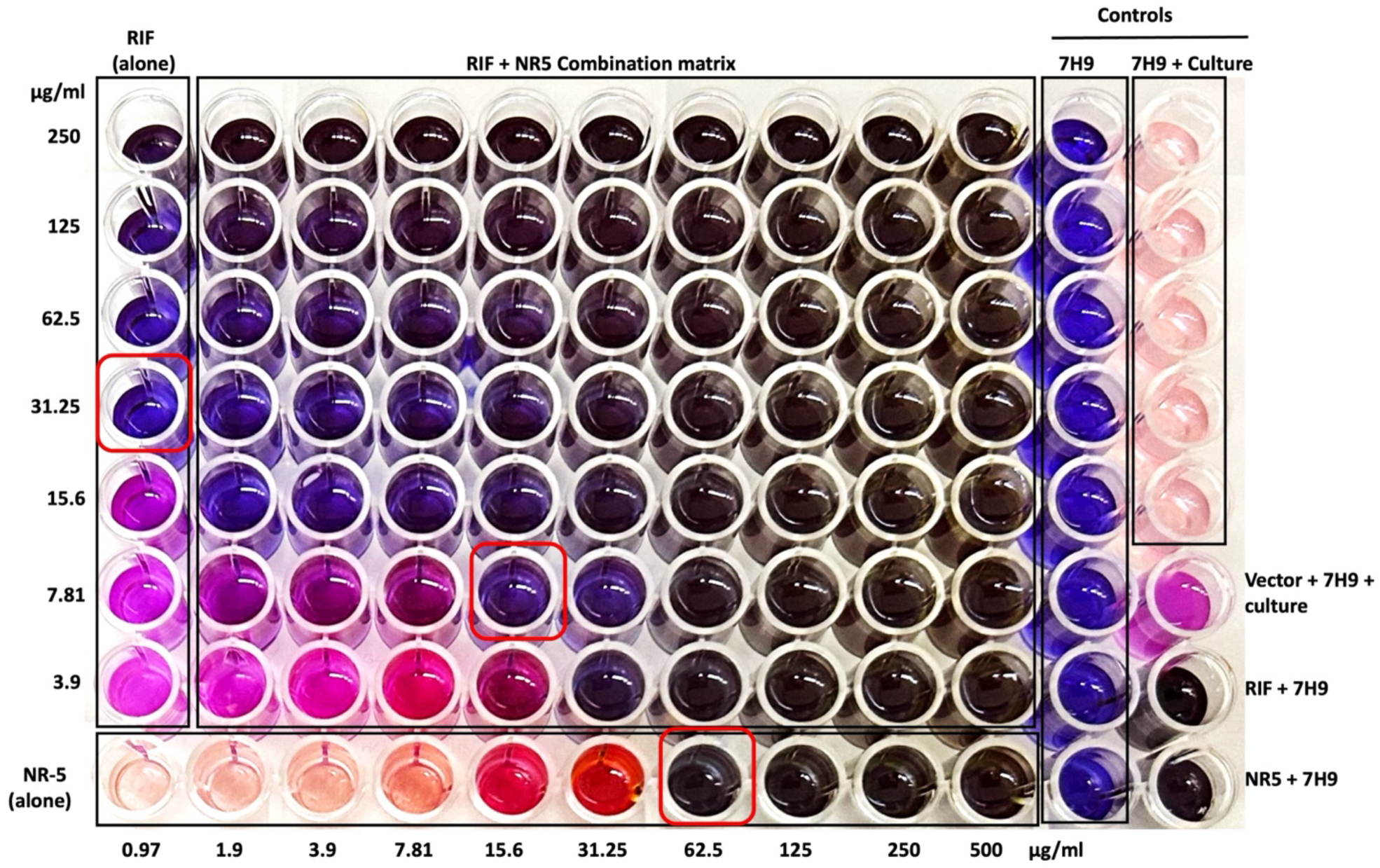
Checkerboard assay evaluating the interaction between NR-5 and rifampicin against *M. smegmatis*. Two-fold serial dilutions of rifampicin (vertical axis) and NR-5 (horizontal axis) were tested in combination. Each well represents a unique concentration pair of the two compounds. The highlighted wells (in red box) indicate the lowest concentration of NR-5 and rifampicin in combination that completely inhibited the growth of *M. smegmatis* and were used to determine the Fractional Inhibitory Concentration Index (FICI). The wells marked with a red box depict the MIC of RIF, NR-5 and the lowest concentration in combination.

**Table 5.** Combination analysis of benzimidazole derivative NR-5 with different first-line and second-line TB drugs against *M. smegmatis*. Combination analysis was qualitatively assessed in three independent experiments (N=3), and consistent interaction patterns were observed across all three experiments.

| S.No. | Antibiotic combinations | MIC (alone)<br>(µg/ml) | MIC (In<br>combination)<br>(µg/ml) | FICI<sup>b</sup> | Type of<br>Interaction |
| --- | --- | --- | --- | --- | --- |
| 01. | Rifampicin<br>+<br>NR-5 | 31.25<br>62.5 | 7.8<br>15.6 | 0.498 | Synergistic |
| 02. | Moxifloxacin<br>+<br>NR-5 | 0.06<br>62.5 | 0.06<br>3.90 | 1.06 | Indifferent |
| 03. | Ethambutol<br>+<br>NR-5 | 0.975<br>62.5 | 0.975<br>62.5 | 2.0 | **Indifferent** |
| 04. | Streptomycin<br>+<br>NR-5 | 0.12<br>62.5 | 0.12<br>31.25 | 1.5 | **Indifferent** |
| 05. | Bedaquiline<br>+<br>NR-5 | 0.012<br>62.5 | 0.012<br>62.5 | 2.0 | **Indifferent** |
| 06. | Kanamycin<br>+<br>NR-5 | 0.0975<br>62.5 | 0.0975<br>31.25 | 1.5 | **Indifferent** |

<sup>b</sup>FICI value of $\leq 0.50$ was defined as synergism, 0.51-1 as additive, 1-4 as indifferent and $>4$ as antagonism.

In contrast, NR-5 demonstrated indifferent interactions with other anti-TB drugs tested, as no significant change in the MIC values was observed in the combination assays. The calculated FICI values ranged from 0.5 to 4 (Table 5), indicating that the antimycobacterial activity is neither enhanced nor antagonised by these drugs.

### 3.8. Molecular Docking Analysis Reveals Potential Targets of NR-5 in *M. smegmatis*

Molecular docking studies of the most active derivative, NR-5, were performed against the 460 essential proteins of *M. smegmatis* to evaluate the binding potential and identify possible molecular targets. The docking analysis demonstrated a range of binding energies across the proteins, highlighting the variation in protein-ligand interactions. Among the proteins tested, NR-5 exhibited the highest binding affinity towards Thymidylate kinase (Tmk), with a docking score of -6.64 Kcal/mol, followed closely by amidophosphoribosyltransferase (PurF) with a docking score of -6.61 Kcal/mol (Table 6). 3′-azido-3′-deoxythymidine monophosphate (AZT) and JNJ-6640 are known inhibitors of Tmk and PurF in *M. tb,* respectively (Fioravanti et al. 2005; Lamprecht et al. 2025), and therefore were taken as positive controls, and ethambutol was taken as a negative control. The docking score of the positive control was -6.99 for AZT and -6.23 for JNJ-6640 (Table 6). Thymidylate kinase (Tmk) is an essential enzyme involved in DNA synthesis. It catalyzes the phosphorylation of thymidine monophosphate (TMP) to thymidine diphosphate (TDP), a critical step in nucleotide metabolism and therefore is necessary for cell viability and proliferation. Due to its role in bacterial survival and its structural differences from the human homolog, Tmk has been recognized as a promising drug target for tuberculosis (Chikhale et al. 2024; Konate et al. 2025). The second-highest scoring target, amidophosphoribosyltransferase (PurF), is involved in de novo synthesis of purines by catalyzing the first committed step in the pathway, which ultimately contributes to DNA synthesis and cell proliferation. The strong binding affinity of NR-5 towards both Tmk and PurF suggests that it may interfere with nucleotide biosynthesis, leading to inhibition of DNA replication and ultimately cell death.

**Table 6.** Molecular docking results of ligands with essential proteins of *Mycobacterium smegmatis*.

| S.No. | Protein | Ligand | Binding Energy (Kcal/mol) |
| --- | --- | --- | --- |
| 1. | Thymidylate kinase (Tmk) | NR-5 | -6.64 |
| 2. | Thymidylate kinase (Tmk) | AZT | -6.99 |
| 3. | Thymidylate kinase (Tmk) | Ethambutol | -4.46 |
| 3. | Amidophosphoribosyltransferase (PurF) | NR-5 | -6.61 |
| 4. | Amidophosphoribosyltransferase (PurF) | JNJ-6640 | -6.23 |

**Table 7.** Molecular interaction profiles of NR-5 and azidothymidine monophosphate (AZT) with thymidylate kinase (Tmk), illustrating non-covalent interactions within the interacting amino acid residues.

| S.No. | Residue | Type of Interaction with NR-5 | Residue | Type of interaction with AZT |
| --- | --- | --- | --- | --- |
| 1 | Asn100 | Hydrogen bond | Asn100, Asp163, Arg74, Asp163 | Hydrogen bond |
| 2 | - | - | Ser99, Tyr103 | Carbon hydrogen bond |
| 3 | | | Arg95 | π- donor hydrogen bond |
| 4 | Phe70, Tyr103 | π-π interaction | - | - |
| 5 | Pro37, Phe70, Met66, Ala67, Val63, Tyr165 | Alkyl, π-alkyl | Pro37 | π-alkyl |

Potential ligand-binding pockets of Tmk and PurF proteins were predicted using the PrankWeb server (Polák et al. 2025). The predicted pockets were ranked based on pocket score, probability, and geometric features. For both Tmk and PurF, the best docking pose of NR-5 was located within the top-ranked predicted binding pocket, supporting the reliability of the docking results.

To further characterize the binding mode of NR-5, protein–ligand interactions were analyzed using PyMOL and Discovery Studio Visualizer. The results indicated that NR-5 occupies the active sites of both proteins and forms multiple interactions with key amino acid residues involved in ligand recognition and stabilization (Fig. 10). In Tmk, NR-5 formed a hydrogen bond with Asn100, π–π stacked interactions with Phe70 and Tyr100, and alkyl or π–alkyl interactions with Pro37, Val63, Met66, Ala67, Arg107, and Tyr165. These interacting residues largely overlapped with those predicted within the binding pocket, validating the relevance of the selected docking region (Fig. 10). Also, the reported Tmk inhibitor, 3′-azido-3′-deoxythymidine monophosphate (AZT) (Fioravanti et al. 2005), was found to occupy the same binding pocket and to share several interacting residues with NR-5 (Fig. 10). The overlap in binding location and interacting residues suggests that NR-5 may targets the catalytic active site of Tmk and may inhibit its enzymatic activity through a mechanism similar to that of known inhibitors.

**Fig. 10.**
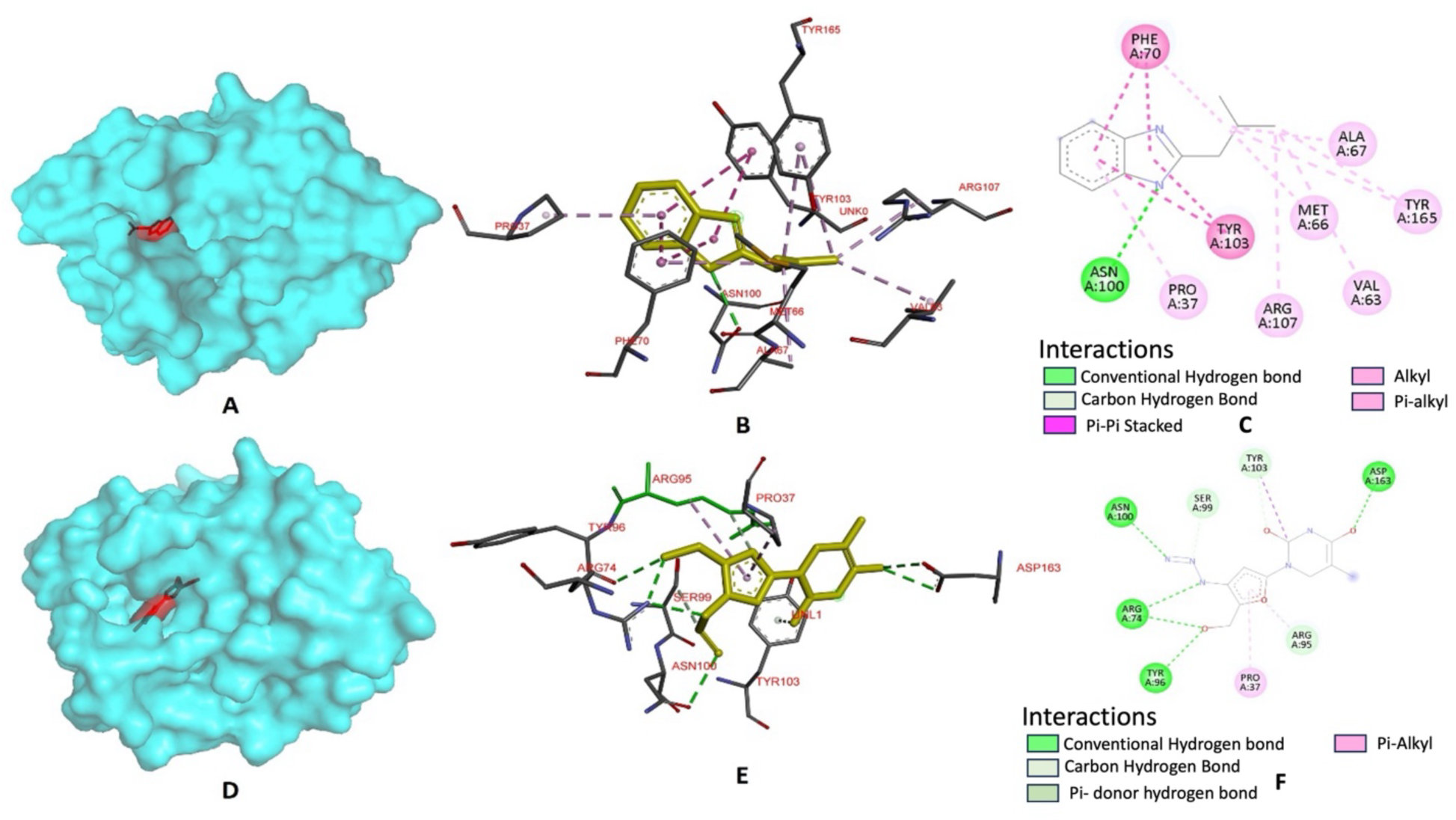
Protein–ligand interaction analysis of NR-5 and AZT with thymidylate kinase (Tmk) of *M. smegmatis*, visualized using PyMOL and Discovery Studio Visualizer. Panels A and D show the surface representation of the binding pocket occupied by NR-5 and AZT, respectively. Panels B and E depict the three-dimensional binding conformations of NR-5 and AZT within the active site of Tmk, highlighting their interactions with key amino acid residues. Panels C and F present the corresponding two-dimensional interaction maps, illustrating hydrogen bonds, π–π stacking interactions, π–alkyl interactions, and π–σ interactions formed between the ligands and surrounding amino acid residues. The overlapping binding location of NR-5 and the known Tmk inhibitor AZT suggests that NR-5 occupies the catalytic active site of the enzyme.

Similarly, in PurF, NR-5 adopted a stable binding conformation within the predicted active pocket through interactions with residues including Asn121, Ala126, Leu480, Gly481, and Val484 (Table 8). The observed interactions, comprising hydrogen bonds, carbon–hydrogen bonds, π–π interactions, and π–σ interactions, are likely to contribute to the stabilization of NR-5 within the binding pocket. Collectively, these findings support the potential of NR-5 to target essential enzymes involved in nucleotide biosynthesis and DNA replication, providing a plausible molecular basis for its observed antimycobacterial activity.

**Table 8.** Identification of ligand-binding pockets of Thymidylate Kinase (Tmk) and Amidophosphoribosyltransferase (PurF) of *M. smegmatis* using the Prankweb server.

| Protein | Ligand | Number of pockets | Pocket based on docking | Pocket Score | Probability | Number of residues | Key Residues |
| --- | --- | --- | --- | --- | --- | --- | --- |
| Thymidylate kinase (Tmk) | NR-5 | 1 | 1 | 25.6 | 0.866 | 30 | Pro37, Val63, Met66, Ala67, Phe70, Asn100, Tyr103, Arg107, Tyr165 |
| Thymidylate Kinase (Tmk) | AZT | 1 | 1 | 25.6 | 0.866 | 30 | Pro37, Arg74, Arg95, Tyr96, Ser99, Asn100, Tyr103, Asp163 |
| Amidophosphoribosyltransferase (PurF) | NR-5 | 8 | 1 | 11.52 | 0.531 | 23 | Thr93, Thr125, Ala479, Leu480, Val123, Asn121, Gly481, Val484, Leu122, Phe405, Thr92, Lys482 |

### 3.9. Structural Stability and Dynamics of the NR-5 – Tmk Complex

The molecular dynamics simulation was performed to investigate the stability of Tmk-NR5 interactions predicted by molecular docking and to evaluate the stability and behaviour of the docked complex. Accordingly, RMSD, RMSF, radius of gyration, solvent-accessible surface area, and hydrogen-bond analyses were performed. From the RMSD plot, it can be observed that the NR-5 RMSD remains relatively high during the early part of the simulation, particularly between 5–30 ns and again around 35–42 ns, where the RMSD is approximately 0.11–0.14 nm (Fig. 11 A). This initial variation is likely due to the ligand adjusting its position within the binding site and undergoing an initial structural rearrangement. After approximately 42 ns, the RMSD gradually becomes more stable and remains mostly within the range of 0.045–0.075 nm (Fig. 11 A). This suggests that the NR-5 adopts a more stable position within the binding pocket during the later part of the simulation. A few occasional sharp spikes can be seen, which likely represent temporary positional or conformational changes. However, these fluctuations are short-lived, and there is no continuous increase in RMSD toward the end of the simulation. Further RMSF analysis revealed that most residues exhibit fluctuations in the range of approximately 0.05–0.12 nm, indicating that the Tmk maintains a relatively stable structure throughout the simulation while still allowing normal residue-level movements (Fig. 11 B). A noticeably higher level of flexibility is observed around residues 155–165, where the RMSF reaches approximately 0.44 nm. This region may correspond to a flexible loop or terminal segment of the protein Tmk and appears to undergo greater movement compared with the rest of the structure (Fig. 11 B). Overall, the protein Tmk remains relatively stable during the simulation, with higher flexibility restricted to a few specific regions rather than showing widespread structural instability. Moreover, it was also observed that the number of hydrogen bonds formed between NR-5 and Tmk throughout the 100 ns simulation (Fig.11 C). The system generally maintains at least one hydrogen bond for a large portion of the simulation. The intermittent loss and re-formation of the hydrogen bond are normal in molecular dynamics simulations and indicate that the interaction is dynamic rather than permanently fixed. Similarly, the minimum distance between Tmk and NR-5 remains relatively consistent throughout the 100 ns simulation, mainly ranging from approximately 0.15–0.21 nm (Fig.11 D). This suggests that NR-5 remains in close contact with Tmk during the simulation, supporting the maintenance of the protein–ligand interaction. Overall, the MD simulation results suggest that the Tmk-NR5 complex maintained relatively stable interactions throughout the 100 ns simulation. These results further support the structural plausibility of the binding interactions predicted by molecular docking, although experimental validation is required to confirm the proposed target.

**Fig. 11.**
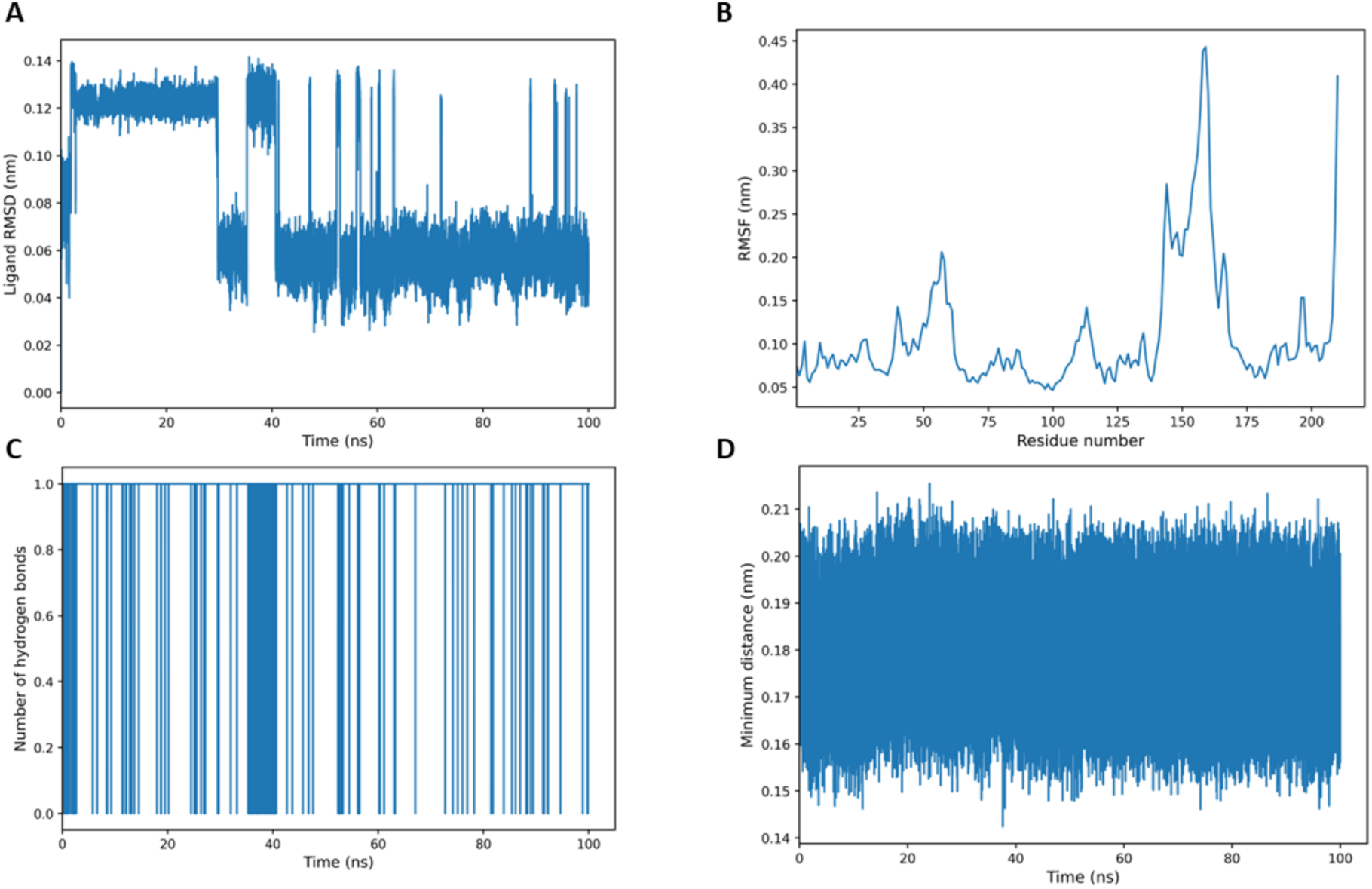
Molecular dynamics analysis of the *M. smegmatis* Tmk-NR5 Complex over 100 ns. (A) RMSD showing changes in the ligand position and stability during the simulation. (B) RMSF indicating the flexibility of individual protein residues. (C) Hydrogen bond analysis showing the formation and breaking of hydrogen bonds between the protein and ligand. (D) Minimum distance between the protein and ligand, reflecting their close interaction throughout the simulation. The plot was generated using a custom script in Python (version 3.13).

## 4. Discussion

The continuous surge in antibiotic resistance and existing therapeutic limitations have intensified the search for novel chemical scaffolds with potent antibacterial activity. Benzimidazole derivatives present as a promising class of compounds due to their broad spectrum of biological activities, including antimicrobial, anticancer, antiviral, and antiparasitic properties (Alheety et al. 2025). In the present study, nine benzimidazole derivatives were evaluated against three bacteria: *M. smegmatis*, *B. subtilis* and *E. coli*. Biological evaluation demonstrated that a set of derivatives (NR-4, NR-5, and NR-7) demonstrated significant antibacterial potency and identified NR-5 as the promising antimycobacterial candidate. However, none of the derivatives was active against *E. coli.* Similar observations have been reported for other benzimidazole-derived compounds, which exhibited limited to no activity against Gram-negative bacteria (Küçükbav et al. 2001; Mohsen et al. 2022).

The resazurin microtitre assay (REMA) showed that, among the tested derivatives, NR-4 and NR-5 inhibited *M. smegmatis growth* with MIC values of 250 μg/mL and 62.5 μg/mL, respectively. However, NR-5 and NR-7 exhibited activity against *B. subtilis* with similar MIC values of 125 μg/mL. Although this MIC value is higher than that of kanamycin (7.8 μg/mL), it is considered acceptable for early-stage lead derivatives and provides a basis for further structural optimizations (Salahuddin et al. 2015; Kantar et al. 2018; Marinescu 2023). Among all derivatives, NR-5 emerged as the most potent antibacterial derivative. Mendogralo et al. (2023) recently reported benzimidazole derivative 3i with an MIC of 7.8 µg/mL and MBC of 62.5 µg/mL against *M. smegmatis*. Although the MIC value of NR-5 is higher than that of 3i against *M. smegmatis*, its MBC value (62.5 µg/mL) is comparable. These findings suggest that NR-5 is an initial antibacterial lead justifying further structural optimization and examination against pathogenic *M. tuberculosis*. The enhanced activity of NR-5 may be attributed to a favourable substitution pattern, electronic effects and lipophilicity, presumably enhancing both cell penetration and target affinity. The comparatively lower MIC value of NR-5 against *M. smegmatis* may suggest a greater affinity for mycobacterial targets as compared to *B. subtilis*. Importantly, NR-5 exhibited bactericidal activity with an MBC/MIC ratio of 1, a desirable characteristic for antimycobacterial agents, as bactericidal compounds are generally preferred for the treatment of mycobacterial infections (Dartois and Dick 2024; Li et al. 2024). The structure–activity relationship (SAR) study of the synthesized benzimidazole derivatives (NR-1–NR-9, Figure 2) uncovered that antibacterial activity is strongly influenced by the electrostatic nature and substitution pattern of chemical groups attached to the benzimidazole core. The compounds without strong electron-modulating groups or containing relatively unsubstituted aromatic scaffolds (NR-1–NR-3 and NR-6–NR-9) displayed negligible antibacterial efficacy, suggesting that the benzimidazole scaffold alone is insufficient for activity. Derivatives NR-4, NR-5, and NR-7 exhibited comparatively higher activity, whereas the remaining compounds showed negligible efficacy. NR-5 demonstrated the most promising activity, potentially due to its favourable electronic properties and lipophilicity.

In addition to antibacterial efficacy, the pharmacokinetic properties of the active derivatives were assessed through in silico ADME analysis. NR-5 demonstrated favourable drug-like properties, in accordance with Lipinski’s rule of five and exhibited high gastrointestinal absorption and oral bioavailability. These observations indicate that these derivatives possess both in vitro efficacy and pharmacokinetic properties that support effective oral administration, thereby supporting their potential for further preclinical studies. However, in silico pharmacokinetic predictions require experimental validation to confirm their physiological relevance.

In contrast, NR-4 violated one physicochemical parameter (insaturation), indicating the need for structural optimization improve its drug-likeness profile. Although NR-7 exhibited antibacterial activity against *B. subtilis*, its increased cytotoxicity toward Vero cells limits its potential as an early hit compound. NR-7, despite showing antibacterial activity against *B. subtilis*, displayed increased cytotoxicity towards Vero cells. NR-7 may be associated with its distinct structural features, particularly the presence of multiple amino groups. These functional groups may influence the compound’s ionization characteristics, interactions with biological macromolecules, and cellular distribution, potentially contributing to its observed cytotoxic profile, consequently minimizing specificity toward bacterial cells. Although substitution improved the antibacterial potency, highly hydrophobic or strongly activating groups might undermine safety margins. The effect of NR-5 on the growth kinetics was also evaluated against *M.* smegmatis. NR-5 significantly inhibited bacterial growth, with complete suppression of growth at 2x MIC and a reduction in growth with a prolonged lag phase at a 1x MIC concentration. It may be hypothesized that this prolongation may be due to inhibition of the biosynthesis of key enzymes required for bacterial metabolism (Rolfe et al. 2012). Pharmacologically, this may reduce the colonization of bacteria at the site of infection at an early stage. Further analysis of time-kill kinetics was also reported, and it was observed that NR-5 gradually reduced the number of viable cells over the period of time. This highlights that NR-5 not only inhibits cell growth but also efficiently kills the bacteria, even under non-growing conditions and in less time, highlighting the effectiveness of NR-5.

Furthermore, combination therapy has emerged as an important strategy, as it enhances the treatment efficacy by targeting multiple cellular pathways simultaneously (Singh et al. 2020). In our study, the checkerboard method was used for combination analysis, and it was observed that NR-5 exhibited synergistic interaction with rifampicin, with a fourfold reduction in the MIC values. The observed synergy may be due to the complementary mechanism of action, as targeting complementary pathways enhances the susceptibility of drugs administered in combination against bacteria (Sullivan et al. 2020). The lack of antagonistic interactions with other drugs used in this study suggests the potential of NR-5 as an antimycobacterial derivative. To further obtain insight into the possible molecular target of NR-5, molecular docking analysis was done against essential proteins of *M. smegmatis*. Among these proteins, thymidylate kinase (Tmk) displayed the highest binding affinity for NR-5, followed by amidophosphoribosyltransferase (PurF). Both these enzymes are key enzymes in nucleotide biosynthesis and DNA replication, making them attractive drug targets. The docking results were further validated through binding pocket prediction and interaction analysis and MD simulation. PrankWeb predicted a single ligand-binding pocket in Tmk, and the best docking pose of NR-5 occupied this pocket, with overlap between the predicted pocket residues and interacting residues. A previous study has also reported the same binding pocket in Tmk, as the predicted residues in this study overlapped with the interacting residues in our study (Hemeda et al. 2023). Notably, the known Tmk inhibitor AZT occupied the same pocket and shared various interacting residues with NR-5. This overlap suggests that NR-5 might occupy the same site of the enzyme. Moreover, it has also been reported that the enzymes involved in thymidine metabolism in *M. smegmatis* are nearly identical to those in *M. tb* and other mycobacterial pathogens (Pecsi et al. 2012). The MD simulation data further supported the docking prediction by demonstrating the relative stability of the Tmk-NR5 complex and consistent intermolecular interactions over 100 ns. These observations suggest a stable predicted binding mode; however, experimental validation is required to establish the biological relevance of the interaction. Moreover, Rifampicin, with which NR-5 have exhibited synergistic activity, inhibits RNA polymerase and thus disrupts RNA synthesis. The docking scores suggest that NR-5 may interact with Tmk and PurF proteins, which are involved in essential metabolic pathways. The simultaneous action of these processes can potentially enhance the antibacterial activity. However, the exact molecular mechanism of the observed synergistic interaction between NR-5 and rifampicin requires further molecular and experimental investigations to identify the mechanism responsible for the enhanced activity.

Overall, the combined microbiological, pharmacological, and computational analyses identify NR-5 as the most promising antimycobacterial benzimidazole derivative evaluated in this study. Its antimycobacterial activity, bactericidal nature, favourable drug-like properties, synergistic interaction with rifampicin, and strong binding affinity toward essential enzymes involved in nucleotide biosynthesis collectively support it as a preliminary antimycobacterial candidate. Consequently, NR-5 is best framed as a promising early-stage hit for further structural optimization and analogue synthesis rather than an immediate clinical therapeutic candidate. Future studies will focus on validating its molecular targets experimentally and also evaluating its efficacy against virulent *Mycobacterium tuberculosis* strains.

## Supporting information

Supplementary Datasheet

## Acknowledgements

Sarika Thakur acknowledges the Department of Biotechnology, India (DBT) for JRF and SRF fellowships. Alka Sharma acknowledges CSIR for Junior Research Fellowship (JRF). Dr Sumitra Nain, Department of Pharmacy, Banasthali Vidyapeeth (for providing benzimidazole derivatives) and Dr Krishna Murari Sinha, Associate Professor, Amity University, Noida (for providing the *M. smegmatis mc^2^ 155* strain) are acknowledged. Dr Ram Gopal Nitharwal is a recipient of the UGC-BSR (F.30-545/2021) and SERB-ANRF (SRG/2020/0001904) grants.

## Author Contributions

Sarika Thakur contributed to data acquisition, experimentation, analysis, as well as drafting and revising the manuscript. Alka Sharma assisted in revising the manuscript. Narayan Kumar helped with molecular dynamics and simulation analysis. Yangala Sudheer Babu and Mulaka Maruthi helped with the experimentation and analysis of the cytotoxicity assay. Mulaka Maruthi, Ravi Kumar, Ram Shankar Upadhayaya and Ram Gopal Nitharwal reviewed and edited the manuscript. Sumitra Nain provided the benzimidazole derivatives and reviewed the manuscript. Ram Gopal Nitharwal supervised the study and guided the research work.

## Supplementary Information

The supplementary information associated with this article is available in the accompanying supplementary files.

## References

1. Abraham MJ, Murtola T, Schulz R, et al (2015) GROMACS: High performance molecular simulations through multi-level parallelism from laptops to supercomputers. SoftwareX 1:19–25. 10.1016/j.softx.2015.06.001

2. Alheety NF, Awad SA, Alheety MA, et al (2025) Benzimidazole Derivatives: A Review of Advances in Synthesis, Biological Potential, Computational Modelling, and Specialized Material Functions. Chemistry (Easton) 8:1. 10.3390/chemistry8010001

3. Amemiya T, Ohkusu K, Murayama M, et al (2024) A rare case of Bacillus subtilis variant natto-induced persistent bacteremia with liver and splenic abscesses in an immunocompetent patient. IDCases 35:e01925. 10.1016/j.idcr.2024.e01925

4. Anichina K, Argirova M, Tzoneva R, et al (2021) 1H-benzimidazole-2-yl hydrazones as tubulin-targeting agents: Synthesis, structural characterization, anthelmintic activity and antiproliferative activity against MCF-7 breast carcinoma cells and molecular docking studies. Chem Biol Interact 345:109540. 10.1016/j.cbi.2021.109540

5. Baindara P, Aslam B (2023) Bacillus spp.-Transmission, pathogenesis, host-pathogen interaction, prevention and treatment. Front. Microbiol. 14:1307723. 10.3389/fmicb.2023.1307723

6. Banerjee P, Kemmler E, Dunkel M, Preissner R (2024) ProTox 3.0: a webserver for the prediction of toxicity of chemicals. Nucleic Acids Res 52:W513–W520. 10.1093/nar/gkae303

7. Banfi E, Scialino G, Monti-Bragadin C (2003) Development of a microdilution method to evaluate Mycobacterium tuberculosis drug susceptibility. J Antimicrob Chemother 52:796–800. 10.1093/jac/dkg439

8. Barák I (2021) Special Issue “Bacillus subtilis as a Model Organism to Study Basic Cell Processes.” Microorganisms 9:2459. 10.3390/microorganisms9122459

9. Baran A, Kwiatkowska A, Potocki L (2023) Antibiotics and bacterial resistance—A short story of an endless arms race. Int J Mol Sci 24:5777. 10.3390/ijms24065777

10. Berenbaum MC (1977) Synergy, additivism and antagonism in immunosuppression. A critical review. Clin Exp Immunol 28:1

11. Bhatt DM, Patel GB, Bhola YO, et al (2026) Hybrids of thiophene, pyrazole and oxadiazol: Synthesis, DFT calculations, molecular docking studies and biological evaluation for anticancer and antimicrobial activity. Synth Commun 56:127–156. 10.1080/00397911.2026.2613352

12. Chauhan M, Rajpoot N, Anthwal T, et al (2025) Synthesis, Characterization and Biological Evaluation of 2-Substituted Benzimidazole Derivatives as Antifungal Compounds. Pharm Chem J 1–6. 10.1007/s11094-025-03387-z

13. Chikhale R V, Pawar SP, Kolpe MS, et al (2024) Identification of mycobacterial Thymidylate kinase inhibitors: a comprehensive pharmacophore, machine learning, molecular docking, and molecular dynamics simulation studies. Mol Divers 28:1947–1964. 10.1007/s11030-024-10967-w

14. Clinical and Laboratory Standards Institute (2023) Performance Standards for Antimicrobial Susceptibility Testing. 33rd ed-M100. Wayne,PA, USA

15. Daina A, Michielin O, Zoete V (2017) SwissADME: a free web tool to evaluate pharmacokinetics, drug-likeness and medicinal chemistry friendliness of small molecules. Sci Rep 7:42717. 10.1038/srep42717

16. Dartois V, Dick T (2024) Therapeutic developments for tuberculosis and nontuberculous mycobacterial lung disease. Nat Rev Drug Discov 23:381–403. 10.1038/s41573-024-00897-5

17. DeJesus MA, Gerrick ER, Xu W, et al (2017) Comprehensive essentiality analysis of the Mycobacterium tuberculosis genome via saturating transposon mutagenesis. mBio 8:10–1128. 10.1128/mbio.02133-16

18. Desai NC, Shihory NR, Kotadiya GM (2014) Facile synthesis of benzimidazole bearing 2-pyridone derivatives as potential antimicrobial agents. Chin Chem Lett 25:305–307. 10.1016/j.cclet.2013.11.026

19. Doern CD (2014) When does 2 plus 2 equal 5? A review of antimicrobial synergy testing. J ClinMicrobiol 52:4124–4128. 10.1128/jcm.01121-14

20. Dragset MS, Ioerger TR, Zhang YJ, et al (2019) Genome-wide phenotypic profiling identifies and categorizes genes required for mycobacterial low iron fitness. Sci Rep 9:11394. 10.1038/s41598-019-47905-y

21. El-Gohary NS, Shaaban MI (2017) Synthesis and biological evaluation of a new series of benzimidazole derivatives as antimicrobial, antiquorum-sensing and antitumor agents. J Med Chem 131:255–262. 10.1016/j.ejmech.2017.03.018

22. Fioravanti E, Adam V, Munier-Lehmann H, Bourgeois D (2005) The crystal structure of Mycobacterium tuberculosis thymidylate kinase in complex with 3 ‘-azidodeoxythymidine monophosphate suggests a mechanism for competitive inhibition. Biochemistry 44:130–137. 10.1021/bi0484163

23. Garcia D, Ramos AJ, Sanchis V, Marin S (2009) Predicting mycotoxins in foods: a review. Food microbiol 26:757–769. 10.1016/j.fm.2009.05.014

24. Hemeda LR, El Hassab MA, Abdelgawad MA, et al (2023) Discovery of pyrimidine-tethered benzothiazole derivatives as novel anti-tubercular agents towards multi-and extensively drug resistant Mycobacterium tuberculosis. J Enzyme Inhib Med Chem 38:2250575. 10.1080/14756366.2023.2250575

25. Herigstad B, Hamilton M, Heersink J (2001) How to optimize the drop plate method for enumerating bacteria. J Microbiol Methods 44:121–129. 10.1016/S0167-7012(00)00241-4

26. Hossain F, Follett P, Vu KD, et al (2016) Evidence for synergistic activity of plant-derived essential oils against fungal pathogens of food. Food Microbiol 53:24–30. 10.1016/j.fm.2015.08.006

27. Ibrahim AA, Said EG, AboulMagd AM, et al (2025) Synthesis and SARs of benzimidazoles: insights into antimicrobial innovation (2018–2024). RSC Adv 15:22097–22127. 10.1039/D5RA00819K

28. Jayavel P, Ramasamy V, Amaladoss N, et al (2024) A facile synthesis, characterization, DFT, ADMET and in-silico molecular docking analysis of novel 4-ethyl acridine-1, 3, 9 (2, 4, 10H)-trione. Chem Phys Impact 8:100476. 10.1016/j.chphi.2024.100476

29. Kantar GK, Mentese E, Beris FS, et al (2018) Synthesis and antimicrobial activity of some new triazole bridged benzimidazole substituted phthalonitrile and phthalocyanines. Rev Roum Chim 63:59–65

30. Khan MT, Zulfiqar T, Jabeen H, et al (2024) Benzimidazole Derivatives in Drug Design: Structure-Activity Relationships and Therapeutic Potential: A Review. J Pharma Bio Med 2:71–80. 10.39401/jpbm.002.02.0018

31. Konate S, Allangba KNPG, Fofana I, et al (2025) Improved inhibitors targeting the thymidylate kinase of multidrug-resistant Mycobacterium tuberculosis with favorable pharmacokinetics. 15:173. 10.3390/life15020173

32. Küçükbav H, Durmaz R, Gueven M, Guenal S (2001) Synthesis of some benzimidazole derivatives and their antibacterial and antifungal activities. Arzneimittelforschung 51:420–424. 10.1055/s-0031-1300057

33. Lamprecht DA, Wall RJ, Leemans A, et al (2025) Targeting de novo purine biosynthesis for tuberculosis treatment. Nature 644:214–220. 10.1038/s41586-025-09177-7

34. Laxminarayan R, Matsoso P, Pant S, et al (2016) Access to effective antimicrobials: a worldwide challenge. Lancet 387:168–175. 10.1016/s0140-6736(15)00474-2

35. Lelovic N, Mitachi K, Yang J, et al (2020) Application of Mycobacterium smegmatis as a surrogate to evaluate drug leads against Mycobacterium tuberculosis. J Antibiot 73:780–789. 10.1038/s41429-020-0320-7

36. Li S-Y, Tyagi S, Soni H, et al (2024) Bactericidal and sterilizing activity of novel regimens combining bedaquiline or TBAJ-587 with GSK2556286 and TBA-7371 in a mouse model of tuberculosis. Antimicrob Agents and Chemother 68:e01562–23. 10.1128/aac.01562-23

37. Marinescu M (2023) Benzimidazole-triazole hybrids as antimicrobial and antiviral agents: A systematic review. Antibiotics 12:1220. 10.3390/antibiotics12071220

38. Miethke M, Pieroni M, Weber T, et al (2021) Towards the sustainable discovery and development of new antibiotics. Nat Rev Chem 5:726–749. 10.1038/s41570-021-00313-1

39. Mohanty SK, Khuntia A, Yellasubbaiah N, et al (2018) Design, synthesis of novel azo derivatives of benzimidazole as potent antibacterial and anti tubercular agents. BJBAS 7:646–651. 10.1016/j.bjbas.2018.07.009

40. Mohsen DH, Radhi AJ, Shaheed DQ, Abbas HK (2022) Synthesis New Benzimidazole Derivatives as Antibacterial. J Pharm Negative Results 13:. 10.47750/pnr.2022.13.S03.137

41. Morcoss MM, Saddik JN, Amin ME, et al (2025) Design, synthesis, antimalarial activity, and in-silico studies of new benzimidazole/pyridine hybrids as dihydrofolate reductase inhibitors. Bioorg Chem 156:108171. 10.1016/j.bioorg.2025.108171

42. Murima P, McKinney JD, Pethe K (2014) Targeting bacterial central metabolism for drug development. Chem Biol 21:1423–1432. 10.1016/j.chembiol.2014.08.020

43. Nepolraj A, Shupeniuk VI, Sathiyaseelan M, Prakash N (2021) Synthesis of new 3-(hydroxymethyl)-2-phenyl-2, 3 dihydroquinolinone and in-silico evaluation of COVID-19 main protease inhibitor. Vietnam J Chem 59:511–521. 10.1002/vjch.202000221

44. Okhuysen PC, Ramesh MS, Louie T, et al (2024) A randomized, double-blind, phase 3 safety and efficacy study of ridinilazole versus vancomycin for treatment of clostridioides difficile infection: clinical outcomes with microbiome and metabolome correlates of response. Clin Infect Dis 78:1462–1472. 10.1093/cid/ciad792

45. Pan T, He X, Chen B, et al (2015) Development of benzimidazole derivatives to inhibit HIV-1 replication through protecting APOBEC3G protein. Eur J Med Chem 95:500–513. 10.1016/j.ejmech.2015.03.050

46. Patel GB, Bhola YO, Dholaria P V, et al (2026) Design, synthesis, and biological evaluation of novel coumarine-1, 2, 3-triazole hybrids as dual DNA gyrase/superoxide dismutase inhibitors: molecular docking, DFT, and ADMET investigations. Future Med Chem 18:1559–1570. 10.1080/17568919.2026.2669609

47. Patel P, Rosani A (2023) Rifampin. In: StatPearls [Internet]. StatPearls Publishing

48. Pathak D V, Maheta JB, Lakhnotra DK, et al (2026) Design, Synthesis, and Anticancer Evaluation of Novel Coumarin-1, 3, 4-Thiadiazole-Chalcone Hybrids as Dual EGFR/VEGFR2 Inhibitors: Molecular Docking, DFT, and ADMET Studies. Polycycl Aromat Compd 1–18. 10.1080/10406638.2026.2642270

49. Pecsi I, Hirmondo R, Brown AC, et al (2012) The dUTPase enzyme is essential in Mycobacterium smegmatis. PLoS One 7:e37461. 10.1371/journal.pone.0037461

50. Polák L, Škoda P, Riedlová K, et al (2025) PrankWeb 4: a modular web server for protein–ligand binding site prediction and downstream analysis. Nucleic acids res 53:W466–W471. 10.1093/nar/gkaf421

51. Rolfe MD, Rice CJ, Lucchini S, et al (2012) Lag phase is a distinct growth phase that prepares bacteria for exponential growth and involves transient metal accumulation. J Bacteriol 194:686–701. 10.1128/JB.06112-11

52. Salahuddin, Mazumder A, Shaharyar M (2015) Synthesis, antibacterial and anticancer evaluation of 5-substituted (1, 3, 4-oxadiazol-2-yl) quinoline. Med Chem Res 24:2514–2528. 10.1007/s00044-014-1308-2

53. Sheikh BA, Bhat BA, Mir MA (2022) Antimicrobial resistance: new insights and therapeutic implications. Appl Microbiol Biotechnol 106:6427–6440. 10.1007/s00253-022-12175-8

54. Singh N, Pandurangan A, Rana K, et al (2012) Benzimidazole: A short review of their antimicrobial activities. Int Curr Pharm J 1:110–118. 10.3329/icpj.v1i5.10284

55. Singh SK, Mohammed A, Alghamdi OA, Husain SM (2020) New approaches for targeting drug resistance through drug combination. In: Combination Therapy Against Multidrug Resistance. Elsevier, pp 221–246. 10.1016/b978-0-12-820576-1.00012-6

56. Song D, Ma S (2016) Recent development of benzimidazole-containing antibacterial agents. Chem Med Chem 11:646–659. 10.1002/cmdc.201600041

57. Stülke J, Grüppen A, Bramkamp M, Pelzer S (2023) Bacillus subtilis, a swiss army knife in science and biotechnology. J Bacteriol 205:e00102–23. 10.1128/jb.00102-23

58. Sullivan GJ, Delgado NN, Maharjan R, Cain AK (2020) How antibiotics work together: molecular mechanisms behind combination therapy. Curr Opin Microbio 57:31–40. 10.1016/j.mib.2020.05.012

59. Tarek A, Jaballah MY, Elrazaz EZ, Samir N (2025) The recent advances in benzimidazole-based antimicrobials and antitubercular agents. Future J Pharm Sci 11:109. 10.1186/s43094-025-00867-7

60. Thakur S, Sharma A, Upadhyaya R, et al (2025) From Bench to Bedside: A Roadmap to Emerging TB Therapeutics. Chem Biol Lett 12:1278. 10.62110/sciencein.cbl.2025.v12.1278

61. Vanommeslaeghe K, Hatcher E, Acharya C, et al (2010) CHARMM general force field: A force field for drug-like molecules compatible with the CHARMM all-atom additive biological force fields. J CompChem 31:671–690. 10.1002/jcc.21367

62. Vianna JF, Bezerra KS, Oliveira JIN, et al (2019) Binding energies of the drugs capreomycin and streptomycin in complex with tuberculosis bacterial ribosome subunits. Phys Chem Chem Phys 21:19192–19200. 10.1039/C9CP03631H

63. Wawrzyniak J (2020) A predictive model for assessment of the risk of mold growth in rapeseeds stored in a bulk as a decision support tool for postharvest management systems. J Am Oil Chem Soc 97:915–927. 10.1002/aocs.12365

64. White RJ, Lancini GC, Silvestri LG (1971) Mechanism of action of rifampin on Mycobacterium smegmatis. J Bacteriol 108:737–741. 10.1128/jb.108.2.737-741.1971

65. WHO (2018) Global antimicrobial resistance surveillance system (GLASS) report: early implementation 2017-2018

66. WHO (2024) Antimicrobial resistance: toolkit for media engagement.

67. Xue F, Luo X, Ye C, et al (2011) Inhibitory properties of 2-substituent-1H-benzimidazole-4-carboxamide derivatives against enteroviruses. Bioorg Med Chem 19:2641–2649. 10.1016/j.bmc.2011.03.007

68. Zwietering MH, Jongenburger I, Rombouts FM, Van’t Riet K (1990) Modeling of the bacterial growth curve. Appl Environ Microbiol 56:1875–1881. 10.1128/aem.56.6.1875-1881.1990

