## Supplementary Tables and Figures for "Identification of a 2-substituted Benzimidazole Derivative with Antimycobacterial Activity and Synergistic Interaction with Rifampicin": Supplementary Information.docx

**Table 1:** Zone of Inhibition against *M. smegmatis*. The experiments were performed independently three times, with each experiment conducted in duplicate

| **Compounds** | **Zone of Inhibition (Mean ± SD)** | | |
| --- | --- | --- | --- |
|  | **Experiment 1** | **Experiment 2** | **Experiment 3** |
| NR-4 | 10.5 ± 0.70 | 11 ± 1.0 | 10.5 ± 0.70 |
| NR-5 | 11 ± 1.41 | 11.5 ± 0.70 | 12 ± 1.41 |
| Rifampicin | 11 ± 0 | 10.5 ± 0.70 | 10.5 ± 0.70 |

**Table 2:** Zone of inhibition against *Bacillus subtilis*. The experiments were performed independently three times, with each experiment conducted in duplicate.

| **Compounds** | **Zone of Inhibition (Mean ± SD)** | | |
| --- | --- | --- | --- |
|  | **Experiment 1** | **Experiment 2** | **Experiment 3** |
| NR-5 | 10.5 ± 0.70 | 12 ± 0 | 10 ± 0 |
| NR-7 | 11.5 ± 0.70 | 11.5 ± 0.70 | 12 ± 0 |
| Kanamycin | 14.5 ± 0.70 | 14 ± 0 | 15 ± 0 |

**Table 3:** Zone of Inhibition (mm) against *M. smegmatis* and *B. subtilis*. The mean value and standard deviation of three experiments performed individually.

| **Bacterial Strains** | **Zone of Inhibition (Mean ± S.D)** | | | | |
| --- | --- | --- | --- | --- | --- |
|  | **NR-4** | **NR-5** | **NR-7** | **Rifampicin** | **Kanamycin** |
| *M. smegmatis* | 10 ± 0.28 | 11.5 ± 0.53 | - | 10.6 ± 0.288 |  |
| *B. subtilis* | - | 10.8 ± 0.84 | 11.63 ± 0.23 | - | 14.53 ± 0.48 |

**Table 4:** In silico prediction of physicochemical properties, pharmacokinetic parameters, drug-likeness, and synthetic accessibility of benzimidazole derivatives (NR-1–NR-9) using SwissADME.

| **Parameter** | **NR-1** | **NR-2** | **NR-3** | **NR-4** | **NR-5** | **NR-6** | **NR-7** | **NR-8** | **NR-9** |
| --- | --- | --- | --- | --- | --- | --- | --- | --- | --- |
| Molecular Weight (g/mol) | 244.25 | 302.50 | 358.60 | 238.28 | 174.24 | 237.51 | 234.3 | 318.41 | 221.26 |
| iLogP | 0.74 | 4.31 | 4.77 | 2.14 | 2.39 | 1.86 | 0.17 | 1.25 | 0.97 |
| TPSA | 84.75 | 24.06 | 24.06 | 41.13 | 24.06 | 24.06 | 111.98 | 44.29 | 87.38 |
| HBA | 3 | 0 | 0 | 1 | 0 | 0 | 2 | 1 | 3 |
| HBD | 5 | 2 | 2 | 2 | 2 | 2 | 6 | 3 | 4 |
| Rotatable Bonds | 1 | 11 | 16 | 2 | 1 | 1 | 5 | 2 | 4 |
| Water solubility | -3.18 | -6.32 | -7.89 | -3.55 | -3.27 | -3.91 | -1.31 | -5.25 | -0.56 |
| Bioavailability score | 0.55 | 0.55 | 0.55 | 0.55 | 0.55 | 0.55 | 0.55 | 0.55 | 0.55 |
| GI absorption | High | High | Low | High | High | High | High | High | High |
| BBB permeability | No | Yes | No | Yes | Yes | Yes | No | Yes | No |
| P-gp substrate prediction | Yes | No | Yes | Yes | No | No | Yes | Yes | Yes |
| Synthetic accessibility | 1.95 | 2.89 | 3.36 | 2.01 | 2.24 | 2 | 2.86 | 2.92 | 2.39 |
| Lipinski violations | 0 | 1 | 1 | 0 | 0 | 0 | 1 | 0 | 0 |
| Veber Violations | 0 | 1 | 1 | 0 | 0 | 0 | 0 | 0 | 0 |

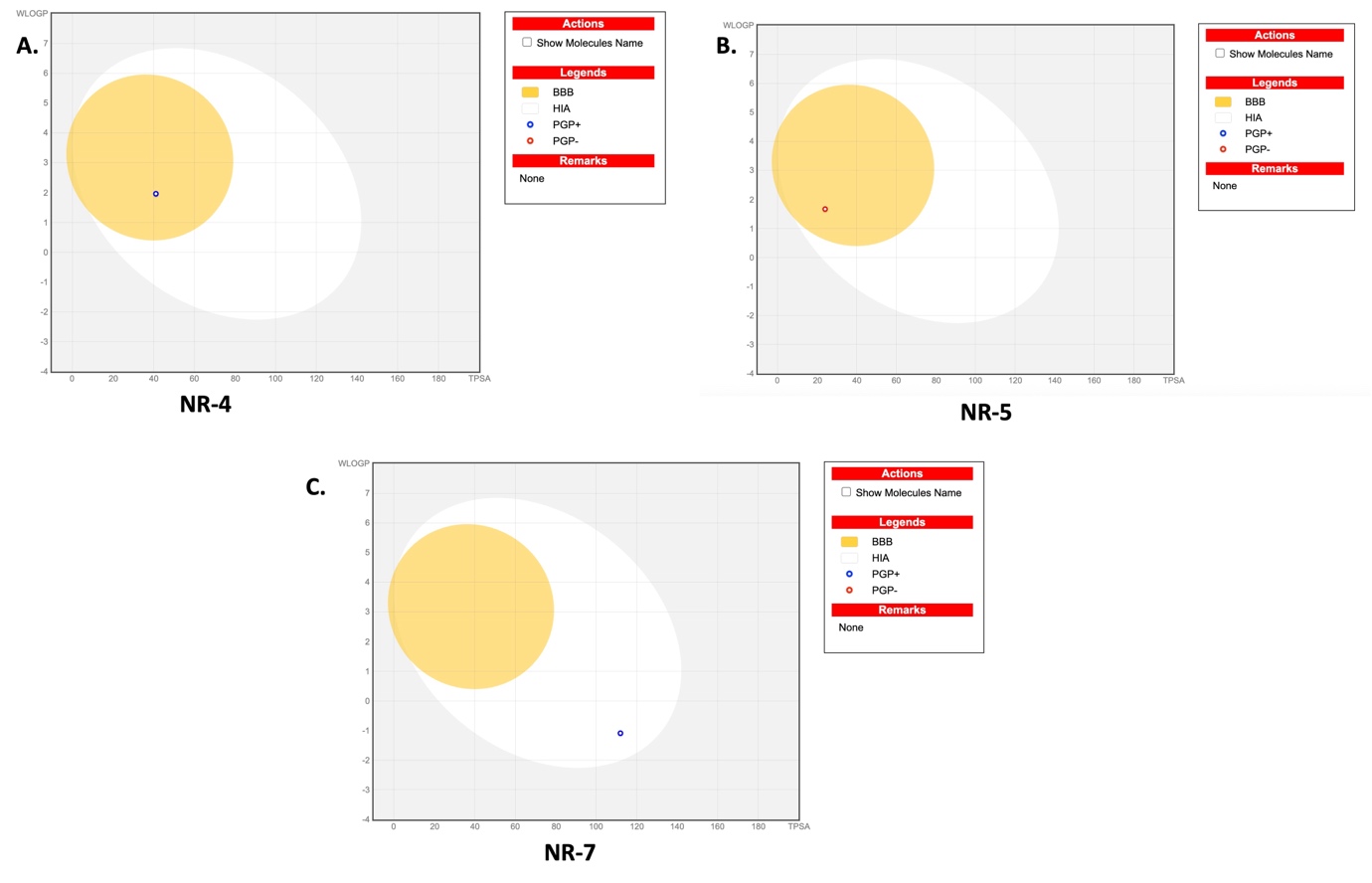

**Supplementary Fig. 1. BOILED-Egg plot of NR-4, NR-5 and NR-7.** BOILED-Egg plot generated using SwissADME to predict gastrointestinal (GI) absorption and blood–brain barrier (BBB) permeability of benzimidazole derivatives. The white region indicates high probability of GI absorption, while the yellow region (yolk) represents high probability of BBB permeation. Blue and red dots indicate predicted P-glycoprotein (P-gp) substrates and non-substrates, respectively.

**Table 5:** In silico toxicity predictions of benzimidazole derivatives (NR-4, NR-5, and NR-7) using ProTox 3.0.

| **Parameter** | **NR-4 (Probability)** | **NR-5 (Probability)** | **NR-7 (Probability)** |
| --- | --- | --- | --- |
| Toxicity Class | 2 | 4 | 5 |
| Hepatotoxicity | Active (0.53) | Inactive (0.57) | Inactive (0.86) |
| Carcinogenicity | Active (0.54) | Inactive (0.55) | Inactive (0.73) |
| Immunotoxicity | Inactive (0.99) | Inactive (0.99) | Inactive (0.73) |
| Mutagenicity | Active (0.53) | Inactive (0.58) | Inactive (0.74) |

**Table 6:** Two-way ANOVA analysis evaluating the effects of treatment, time, and their interaction on bacterial growth, showing the percentage contribution to total variation and corresponding statistical significance (***P < 0.0001). All three factors exhibited statistically significant effects.

| **Source of Variation** | **% of total variation** | **P value** |
| --- | --- | --- |
| Interaction | 27.13 | P<0.0001 |
| Time (hours) | 23.65 | P<0.0001 |
| Treatment | 49.05 | P<0.0001 |
| Source of Variation | P value summary | Significant? |
| Interaction | *** | Yes |
| Time (hours) | *** | Yes |
| Treatment | *** | Yes |
